# A systems-biology approach to molecular machines: Exploration of alternative transporter mechanisms

**DOI:** 10.1101/685933

**Authors:** August George, Paola Bisignano, John M. Rosenberg, Michael Grabe, Daniel M. Zuckerman

**Affiliations:** Dept. of Biomedical Engineering, Oregon Health and Science University, Portland, OR, USA; Dept. of Pharm. Chem., University of California San Francisco, San Francisco, CA, USA; Dept. of Biological Sciences, University of Pittsburgh, Pittsburgh, PA, USA

## Abstract

Motivated by growing evidence for pathway heterogeneity and alternative functions of molecular machines, we demonstrate a computational approach for investigating two questions: (1) Are there multiple mechanisms (state-space pathways) by which a machine can perform a given function, such as cotransport across a membrane? (2) How can additional functionality, such as proofreading/error-correction, be built into machine function using standard biochemical processes? Answers to these questions will aid both the understanding of molecular-scale cell biology and the design of synthetic machines. Focusing on transport in this initial study, we sample a variety of mechanisms by employing Metropolis Markov chain Monte Carlo. Trial moves adjust transition rates among an automatically generated set of conformational and binding states while maintaining fidelity to thermodynamic principles and a user-supplied fitness/functionality goal. Each accepted move generates a new model. The simulations yield both single and mixed reaction pathways for cotransport in a simple environment with a single substrate along with a driving ion. In a “competitive” environment including an additional decoy substrate, several qualitatively distinct reaction pathways are found which are capable of extremely high discrimination coupled to a leak of the driving ion, akin to proofreading. The array of functional models would be difficult to find by intuition alone in the complex state-spaces of interest.

**Author summary:** Molecular machines, which operate on the nanoscale, are proteins/complexes that perform remarkable tasks such as the selective absorption of nutrients into the cell by transporters. These complex machines are often described using a fairly simple set of states and transitions that may not account for the stochasticity and heterogeneity generally expected at the nanoscale at body temperature. New tools are needed to study the full array of possibilities. This study presents a novel in silico method to systematically generate testable molecular-machine kinetic models and explore alternative mechanisms, applied first to membrane transport proteins. Our initial results suggest these transport machines may contain mechanisms which ‘detoxify’ the cell of an unwanted toxin, as well as significantly discriminate against the import of the toxin. This novel approach should aid the experimental study of key physiological processes such as renal glucose re-absorption, rational drug design, and potentially the development of synthetic machines.

## Introduction

The proteins and protein complexes known as molecular machines perform essential functions in the cell, including transport, locomotion, energy production, and gene expression [1]. Secondary active transporters, the focus of the present study, move ions and small molecules across a membrane driven by an electrochemical gradient of an ion [1]. For example, SGLT transporters are of biomedical interest due to the vital role that SGLT1 and SGLT2 play in the uptake of glucose in the small intestines and reabsorption in the kidneys, respectively [2], which in turn has prompted biophysical scrutiny of their mechanisms [3–7]. Numerous other transporters have also been assayed on a quantitative basis [8–13].

The biological mechanisms of transporters as well as other molecular machines can be modeled using chemical reaction networks, typically along with mass action kinetics [14]. In a chemical reaction network, the system process is decomposed into discrete states connected by transition rates between states [14] forming a network of interconnected reactions in the state-space (Fig. 1). These networks can be modeled using the chemical master equation: a set of differential equations describing the state probabilities and connected transition rates for each state [15]. Biochemical networks are generally Markovian [16], have a number of different control patterns [17], and typically adhere to specific design principles [18].

**Fig 1.**
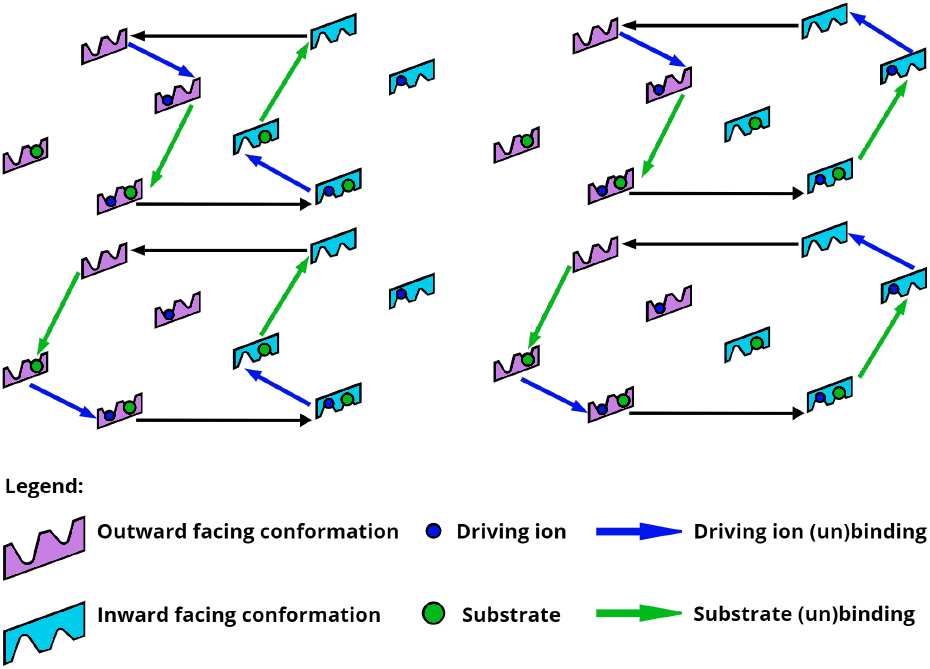
Multiple mechanisms for ideal symport. The four “ideal” kinetic pathways of a hypothetical symporter that transports substrate using the available free energy of the driving ion are shown. This state-space contains eight states, and symport models include at least six connecting transitions.

Despite the complex state-spaces accessible to molecular machines such as transporters, their mechanisms are often described using single-pathway, highly machine-like cartoon-like models [1, 19, 20], building on the seminal suggestions of Mitchell [21] and Jardetzky [22]. See Fig. 1. While such models are helpful for a qualitative understanding of complex protein behavior and chemical networks, simple models may also build in unwarranted assumptions about the system. There is growing evidence that molecular machines may exhibit complexity beyond that embodied in typical ‘textbook’ cartoons models [8, 9]. Recent experimental studies have shown that certain traditional model assumptions such as fixed stoichiometry [10, 11], homogeneous pathways [12], and unique binding sites [13] may be incorrect.

As an example of mechanistic alternatives within a simple state-space, consider a hypothetical cotransporter motivated by the SGLT symporter which transports a single substrate and is driven by an ion gradient. The state-space is constructed using three state ‘dimensions’: conformational state, ion binding state, and substrate binding state. For this hypothetical transporter there are two conformations, each permitting four ion/substrate binding states: fully unbound, ion bound, substrate bound, and fully bound. Within this relatively simple state-space, we can construct four ideal kinetic pathways (**Fig. 1**) that connect the minimum number of states to produce a symport cycle (i.e., intracellular transport of the substrate coupled to ion flow). However, there are numerous additional mechanistic possibilities: combinations of the ideal pathways, or even non-ideal pathways including, e.g., an ion leak. Note that antiport cycles can be similarly constructed using this same state-space [23].

Beyond the nominal functions of transporters, we also take note of one of the most remarkable properties of some molecular machines: the ability to perform “proofreading” or error-correction [24–26]. Specifically, certain network topologies promote enhanced selectivity (i.e., reduced error) in systems with a competing substrate [24–26]; this enhancement in selectivity incurs a free energy cost, typically paid via hydrolysis of a phosphodiester bond. While some aspects of proofreading networks have been examined – such as the speed, accuracy, and dissipation trade-offs [27, 28], as well as non-equilibrium proofreading regimes [29] – the possibility that *transporters* might exhibit proofreading has not been explored to our knowledge.

Here we pursue a systematic exploration of mechanistic and functional diversity, building on strategies developed largely within the field of systems biology. Due to the challenges of modeling complex biochemical networks, such as enumerating combinatorically large state-spaces, new approaches have been developed to keep these systems tractable. Genetic-algorithm sampling has been used to evolve complex biochemical networks such as metabolic pathways [30, 31]. In related work applied to ion channels, models have been fit to experimental data using genetic algorithms and simulated annealing [32–34]. These reverse-engineering studies primarily sought optimal individual models, not the model-sets we pursue here, and also did not account for non-equilibrium constraints on transition rates [14].

Motivated by the possibility of discovering new potential mechanisms and the limitations of current “manual” approaches for analyzing molecular machines, we systematically explore the biochemical network model space for a given molecular machine or function. Our approach generates diverse models that are both thermodynamically consistent and testable. Due to their biomedical significance, we first examined transporters motivated by SGLT-type proteins. Our results suggest there is a diverse set of possible mechanisms for these cotransporters, including proofreading driven by an ion leak.

## Methods

We have developed custom software, ModelExplorer, to study molecular machine behavior. Although this study is focused primarily on membrane cotransporters, the software is designed to be general enough for the exploration of other molecular systems. ModelExplorer automatically generates combinatorial state-spaces and then uses a modified Monte Carlo Metropolis algorithm to sample the model space with a user-defined “energy” or fitness function. This fitness function could embody experimental measurements via a weighted sum of residuals (loss function), but here we use functionality-motivated fitness functions. The software can also impose constraints motivated by structural or biochemical knowledge, such as prohibited states or a known order of binding events.

### Model specification

We create a system model consisting of states and connecting rate constants. Systems states are created from all the allowed combinations of user-specified conformational and chemical substates, and placed into physically equivalent groupings (**see SI-1**). The Monte Carlo sampling generates a trajectory in model space (Fig. 2 and see below) that allows the selection of the fittest models, with a tempering procedure used to avoid trapping. Each model is assessed by its steady-state behavior in the current implementation, although transient information could be employed. The generated models may then be analyzed for kinetic pathways as well as flow stoichiometry over a range of chemical potential conditions, resulting in experimentally testable models.

**Fig 2.**
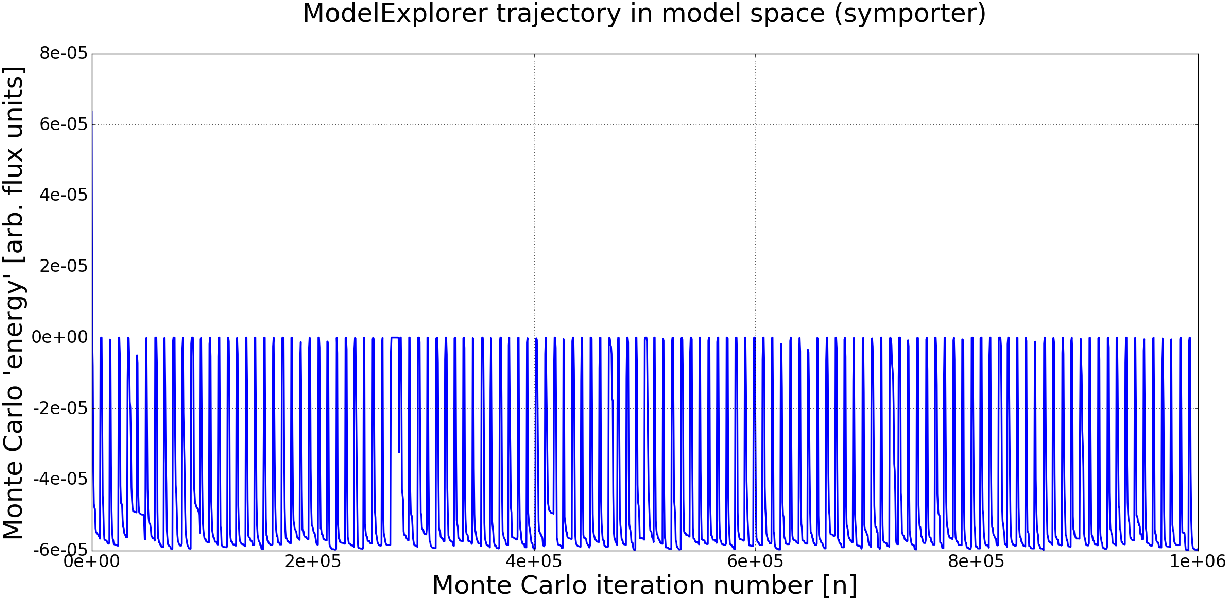
Exploration of model space using Markov-chain Monte Carlo. The plot shows a ModelExplorer trajectory based on symporter energy function during a 1e6 MC step simulation. Note that each point represents a different fully specified model. Energy minima correspond to models that are more fit, using *E*_MC_ = −*J*_substrate_ as a fitness function in this case, where *J*_substrate_ is the flux of substrate. Models are initially at a high energy but quickly find local minima. A tempering schedule of alternating temperature increases and decreases prevents the simulation from being trapped in local minima.

The behavior of each model is determined by the rate constants governing transitions among the states. To ensure thermodynamic consistency among all rate constants [14], we use an energy-based formulation [35–37] where a free energy value is assigned to each state and transition state. To simplify equations, we use reduced units, with all energies expressed in units of *k_B_T*. For a conformational transition from state *i* → *j*, the Arrhenius-like first-order rate constant is expressed using Hill’s notation [14] as:

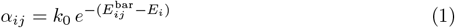

The user-defined prefactor *k*_0_ is set arbitrarily to 10^−3^ *s*^−1^, while 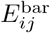 corresponds to the transition state (free) energy and 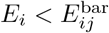 is the free energy of state *i*. Because *k*_0_ is the pre-factor for all rate constants, its value does not affect the mechanisms discovered, but rather it influences the overall rates as a scaling factor.

For a process *i* → *j* involving binding or unbinding, the ion or substrate concentration is built into an effective first-order rate constant [14] given by:

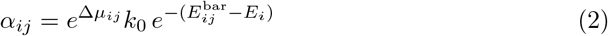

with the same parameters as above, and with Δ*μ_ij_* being the non-equilibrium difference in the chemical potential for the *i* → *j* state transition:

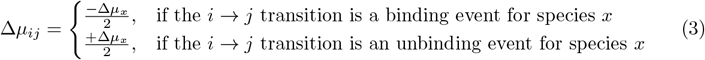

Here, Δ*μ_x_* is the difference in the chemical potential across the membrane for species *x* (e.g. ion or substrate) [14]:

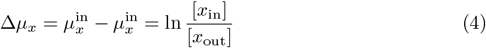

where [*x*_in_] and [*x*_out_] are the intracellular and extracellular concentrations of species *x*. Note that by construction only one species can have a binding/unbinding event during an *i* → *j* state transition. The factor of 1/2 in eq. 3 is an arbitrary choice for dividing the driving force and was found empirically to have a minimal effect on the models found. Note that eq. 4 does not include the membrane potential for charged species (e.g., a sodium ion), which is excluded from this study to focus on the simplest cases.

We use a novel string-based approach to construct the state-space for a molecular machine in a combinatorial fashion, filtering out user-defined exclusions. This is best illustrated with an example for a hypothetical alternating-access transporter of substrate (S) driven by a sodium ion (N): the state “OF-Nb-Si” represents the outward-facing (OF) conformation with a sodium bound (Nb) and substrate inside the pertinent cell or organelle. The string OF-Nb-Si fully defines the system state in a sufficient way for our kinetic scheme explained below. The state string consists of a set of user-defined ‘base state’ strings, each of which describes a different characteristic of the machine. For a transporter, this includes the conformational state (inward or outward facing; IF or OF), sodium-ion state (bound to protein, intracellular/”inside”, or extracellular/”outside”; Nb, Ni, or No), and a substrate binding state (Sb, Si, or So), resulting in a 3D state-space. The inside and outside designations are necessary to define the direction of a transition (e.g., N binding from outside) but note that under the steady-state conditions analyzed here, species concentrations are held fixed but with differing values inside and outside. That is, transitions do not change the steady-state concentrations.

To examine transport mechanisms in heterogeneous environments, we add an additional binding base state for the competing substrate (W) following the same conventions (Wb, Wi, or Wo). These states are defined in an analogous manner: e.g., OF-Nb-Wi. By choice, S and W cannot both be bound simultaneously.

Transitions are assumed to occur only via single elementary steps, such as a single (un)binding event or conformational change. Transitions between states that involve more than one conformational or binding state change are not permitted. Transition rate constants are defined using an Arrhenius scheme as noted above in equations 1 and 2. Ultimately, steady-state populations are calculated for distinguishable states. For example, the “OF-Nb-Si” and “OF-Nb-So” states are not distinguishable in steady state because inside and outside substrate concentrations are held fixed. (As noted above, the inside and outside designations are necessary to assign directionality to binding events.) In contrast, the “OF-Nb-Si” and “OF-Ni-Sb” states differ in their binding state making them distinguishable. Likewise the “OF-Nb-Si” and “IF-Nb-Si” states are distinguishable. Hence, as detailed in the SI, the full set of states is mapped to a smaller set of physically unique states for determining populations. Furthermore, any set of indistinguishable states must be energetically equivalent with the same *E_i_* value in equations 1 and 2. These indistinguishable states are “tied” together for the Monte Carlo procedure. Note that state populations cannot be directly inferred from the equilibrium *E_i_* values because we are studying driven, non-equilibrium steady states.

### Model sampling

We use the Monte Carlo (MC) Metropolis-Hastings algorithm [38, 39] with tempering to sample model space, where a trial move consists of randomly adjusting a single state or transition energy, *E_i_* or *E_ij_*, along with the energies of its equivalent tied members. As noted above, the tied members consist of states which are physically equivalent given a steady state where the outside and inside concentrations are fixed. Physically equivalent transitions are those occurring between physically equivalent states; see SI. Adjusting all the “tied” states during a trial move prevents physically equivalent states in the model from having different free energies, maintaining thermodynamic consistency.

For each trial model generated using MC, we evaluate its MC “energy” or fitness based on its steady-state characteristics. To do so, we construct a rate matrix, K, for the new model using the equilibrium and non-equilibrium (binding) energies for each state/transition along with the modified Arrhenius equations, (eq. 1 and 2). The rate matrix, **K**, can be written with the state probabilities column vector, 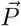, to form the chemical master equation [15]:

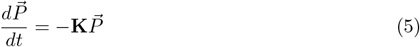

where *K_ij_* = −*α_ji_* for each transition with *i* ≠ *j*, and *K_ii_* is the sum over outgoing rate constants, ∑_*j*≠*i*_ *α_ij_*. The chemical master equation can be solved under steady-state conditions using the definition of steady-state, 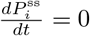, and the additional constraint that ∑*_i_ P_i_* = 1. This yields the steady-state probabilities for each state, from which we can calculate the steady flows between states when combined with the rates [40]:

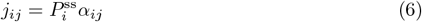

where *j_ij_* is the steady-state flow from state *i* to state *j*, 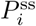 is the steady-state probability of being in state *i*, and *α_ij_* is the transition rate between state *i* and *j*. The overall flux, *J_x_*, of a species (e.g. substrate or ion) is then determined from the sum of net flows of user defined transitions, such as the substrate binding/unbinding transitions in the ‘inward’ facing conformation (see SI):

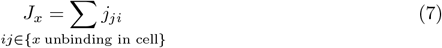

Each model (set of rate constants between states) will have a Monte Carlo ‘energy’ *E_MC_* assigned (i.e. fitness score) based on a user-defined general function. In this study we use substrate flows into the cell, as specified in the Results section for different choices of the fitness function.

Each Monte Carlo step results in a model, and thus each simulation yields a trajectory in “model space” (**Fig. 2**) – i.e., a sequence of models. By convention, lower fitness scores are more fit. The current model’s ‘energy’ is compared to the previous model’s energy and accepted/rejected based on the Metropolis-Hastings selection criterion [38, 39].

That is, the probability of accepting a trial move is the usual *p* = min[1, *e*^−*β*Δ*E_MC_*^], where Δ*E*_MC_ is the change in the Monte Carlo ‘energies’ of the two models, and *β* is the effective inverse thermal-energy parameter. We emphasize, however, that our MC procedure does not generate a true Boltzmann-distributed thermal ensemble, and the MC *β* parameter is used only to aid sampling.

In order to avoid trapping in deep “energy” (high fitness) basins we employ a tempering [41] procedure. This procedure cyclically raises and lowers the inverse-temperature *β* in the Metropolis-Hastings acceptance criterion, facilitating the exploration of different areas in the model space because of the increased likelihood of acceptance. ModelExplorer allows for both adaptive and fixed-cycle tempering. Adaptive tempering tracks the change in fitness of the previous models, decreasing *β* (heating) if the fitness has not changed over several models, and then increasing *β* (cooling) once a user-defined threshold is met (see SI). Fixed-cycle tempering sets a fixed heating and cooling schedule for the duration of the simulation. The tempering procedure is fully customizable and increases the diversity of models found in a simulation. Figure 2 is an example of the Monte Carlo “energy” trajectory in model space produced in ModelExplorer for a simple symporter system using fixed-cycle tempering. Note that the initial models which are not easily visible at the very left of the graph exhibit poor fitness (positive MC energy), but negative-energy models are quickly found.

### Model analysis

Models can be analyzed in several ways using ModelExplorer: characterizing overall stoichiometries, kinetic pathway analysis, and manual adjustment of selected transition rates. The ion and substrate flux of a model can be calculated over a range of chemical potential differences by incrementing the desired chemical potential difference and updating the non-equilibrium energy terms for each state. The rates, steady state probabilities, flows, and then fluxes are then recalculated for each chemical potential increment, allowing for the examination of stoichiometry. Since ModelExplorer calculates the net flows between states, a kinetic diagram of the network pathways can be made for a model at user-defined chemical potential differences. Note that the absolute flow (and flux) values are not physically relevant due to an arbitrary overall rate constant prefactor: all flows can be scaled by an arbitrary constant and remain consonant with the governing equations.

Furthermore, individual models may be perturbed by modifying specific state/transition energies, leading to insights on pathway characteristics (e.g. pathway with or without leaks). Combining the kinetic pathway diagram with the flux analysis (**Fig. 3**) provides a means to investigate the dynamic behavior of molecular machines and provide testable mechanistic hypotheses. Since the flow and flux values are arbitrary (see above) the ion and substrate flux have been scaled by the maximum flux (i.e. ion flux at largest chemical potential difference).

**Fig 3.**
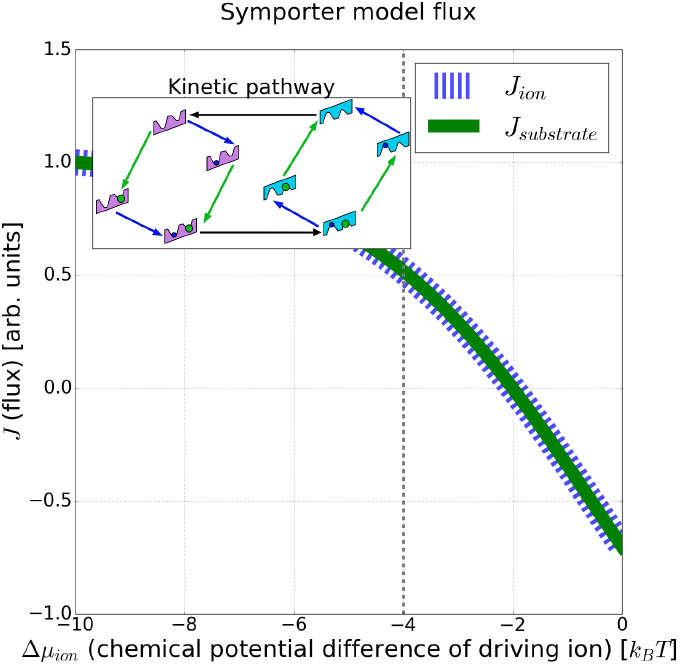
Ideal symport in a non-ideal (mixed) model. The fluxes, *J*, of the driving ion and substrate of an example symporter model are plotted over a range of ion chemical potential differences. Note the 1:1 stoichiometry of the substrate to ion flux, indicating an ideal symporter with no leaks. The ion and substrate flux have been scaled by the maximum ion flux for visual clarity. Inset: kinetic pathway of the same symporter model at an ion chemical potential difference of −4*k_B_T*, as indicated by the vertical line.

In addition to single model analysis, we have developed tools for the meta-analysis of the data – i.e., a “systems” analysis. A simple data pipeline allows for the analysis of sampling parameters, run-to-run model distributions, and model clustering, based on the differences in model flows. Sampling efficiency can be evaluated based on the average time to find a sufficiently different model (i.e. cluster) during the simulation as well as the run-to-run comparison. The run-to-run comparison tool computes the minimum distance between models found in simulation ‘A’ compared to simulation ‘B’ and vice versa, generating a model distribution between the runs. This provides insight into unique models found under different simulation conditions. Models may also be clustered hierarchically [42], allowing for the discovery of different model classes. The model differences are calculated using the Euclidean distance between the vectors of the scaled (by maximum) net flows between the states of a given model.

ModelExplorer has a number of adjustable physical parameters (change in chemical potential), simulation parameters (seed, run length), sampling parameters (tempering schedule, amount to perturb a state/barrier energy), as well as constraints (illegal transitions and states) for a simple cotransport system. In order to investigate Hopfield-like enhanced discrimination [24], the cotransport system with a decoy substrate has additional parameters and constraints. The decoy-bound state energies differ from their equivalent substrate-bound states by a fixed amount (ΔΔ*G* parameter), and have equal barrier energies. This constraint forces the enhanced selectivity to result from the difference in binding affinities alone (ΔΔ*G*), and not “internal proofreading”, which is consistent with Hopfield’s kinetic proofreading network [24]. See Discussion. We noted that occluded states (open to neither inside nor outside) are omitted for simplicity, and that decoy and substrate are mutually exclusive binders.

### Computing details

The current prototype of ModelExplorer was written in Perl 5 with additional scripts for analysis and visualization written using Python 2.7. The software was used on a desktop PC running Windows 10 64-bit OS with an Intel i7-6700 CPU. Each simulation was run for 1e6 MC steps, storing models every 500 MC steps to reduce the run-time and memory requirements. Each run yielded 2000 models. For simulation parameters **see SI-2**. All scripts and data for the manuscript are available at: https://github.com/ZuckermanLab/ModelExplorer.

## Results

We present results for a range of systems demonstrating the ability of the computational approach to discover both expected and surprising mechanisms in single and competing-substrate environments. For a simple cotransporter system without a decoy, the results include validation of ideal symporter/antiporter mechanisms, selected network topologies, and substrate/ion fluxes for a range of ion chemical potential differences. For cotransporters with an additional decoy substrate, we analyze several features of four simulations: selected network topologies, substrate/decoy/ion flux for varying ion chemical potential differences, network heterogeneity via clustering, and possible alternative mechanisms.

### Ideal and mixed-mechanism cotransport with a single substrate

We first studied a transporter system with a single substrate driven by an ion gradient. To generate symport models, the substrate chemical potential difference (Δ*μ*_substrate_) was set to 2*k_B_T* and the ion chemical potential difference (Δ*μ*_ion_) was set to −4*k_B_T*; see (4). The fitness goal as embodied in the Monte Carlo energy was set to:

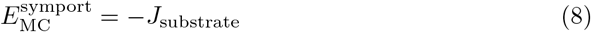

where *J*_substrate_ is the flux of the substrate into the cell as given by (7). This promotes intracellular substrate flow against its own gradient bud down the ion gradient. Note that negative Monte Carlo energies are better by convention. The simulation of 1e6 MC steps yielded the four idealized symporter cycles of Fig. 1, as well as combinations of the idealized symporter cycles (Fig. 3 and **see SI-3**). All of these models exhibit a 1:1 ratio of substrate and ion flux into the cell – consistent with the idealized predictions of an optimized symporter (Fig. 1).

Antiporter behavior in the same state-space was explored by modifying the substrate chemical potential difference (Δ*μ*_substrate_) to −2*k_B_T* and setting the fitness goal to:

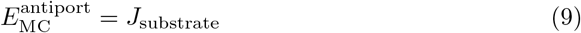

where *J*_substrate_ is the flux of the substrate. Because MC favors lower energy, the use of (9) promotes extracellular flow of the substrate opposite to the ion gradient and flow. The simulation of 1e6 MC steps yielded antiporter models with a 1:1 ratio of substrate flux out of the cell to ion flux into the cell (see SI) – consistent with theoretical expectations for an optimized antiporter.

### Discriminative models in the presence of a competing substrate

A primary motivation for this work was the challenge of generating models with particular functions in complex state-spaces where a combinatorial number of possibilities preclude guessing of mechanisms based on intuition. Specifically, motivated by hints that vSGLT exhibited non-productive reversal events (substrate unbinding to the extracellular side) [43], we wanted to investigate whether slippage events [14] might be able to enhance selectivity in the presence of a “decoy” substrate.

To seek models capable of enhancing selectivity for one substrate over another (*beyond* that generated by their differing affinities, importantly), a competing decoy substrate was added to a transport state-space that included a driving ion gradient. The decoy substrate was set to have a weaker affinity by ΔΔ*G* = 1*k_B_T*, but otherwise the substrates are treated identically. The substrate and decoy chemical potential differences (Δ*μ*_substrate_, Δ*μ*_decoy_) were both set to 2*k_B_T*, and the ion chemical potential difference (Δ*μ*_ion_) was set to −4*k_B_T*. The fitness function (Monte Carlo energy) was set to:

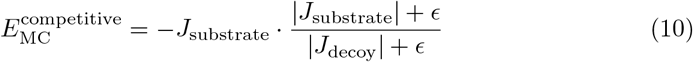

where, *ϵ* = 1e–15 improves numerical stability, and *J*_substrate_ and *J*_decoy_ are the flux of the substrate and decoy respectively as defined in (7). The first factor of equation 10 promotes the intracellular flux of the substrate (negative sign used by convention), while the second factor promotes a high ratio of substrate to decoy flux (i.e. enhanced selectivity). Note that while the models are optimized at an ion chemical potential difference of −4*k_B_T*, we are also able to find enhanced discrimination at larger ion chemical potential difference (i.e. concentration) values.

### Analysis of a single discriminative model

To highlight the power of the sampling strategy to discover non-trivial mechanisms, we examine the kinetic pathways of a model (**Fig. 4A**) with enhanced selectivity. (Below this is referred to as “Model B” – MC index 29000.1, **see SI-4**). This model can be described as a combination of several pathways: two pathways which symport both the ion and substrate (Fig. 4C), and two ion leak pathways which only transport the ion (Fig. 4B, D). Here we have defined an ion leak as a “futile” cycle in the state-space leading solely to dissipation of the ion gradient. We can intuitively understand the discrimination mechanism: the ion leak pathways (which include the central horizontal arrow in Figure 4B, D) drive the substrate and decoy to unbind in the outward facing conformation (on the left side of the diagrams). Due to the 1*k_B_T* difference in binding affinities, the substrate is more likely to rebind and be transported into the intracellular region. In fact, under the conditions which lead to the flows shown in Figure 4A, there is negligible decoy flow into the cell, which is why no flow arrow is shown connecting the two decoy-and-ion bound states. The mechanism of this model directly echoes the driven tRNA unbinding from the ribosome analyzed by Hopfield [24].

**Fig 4.**
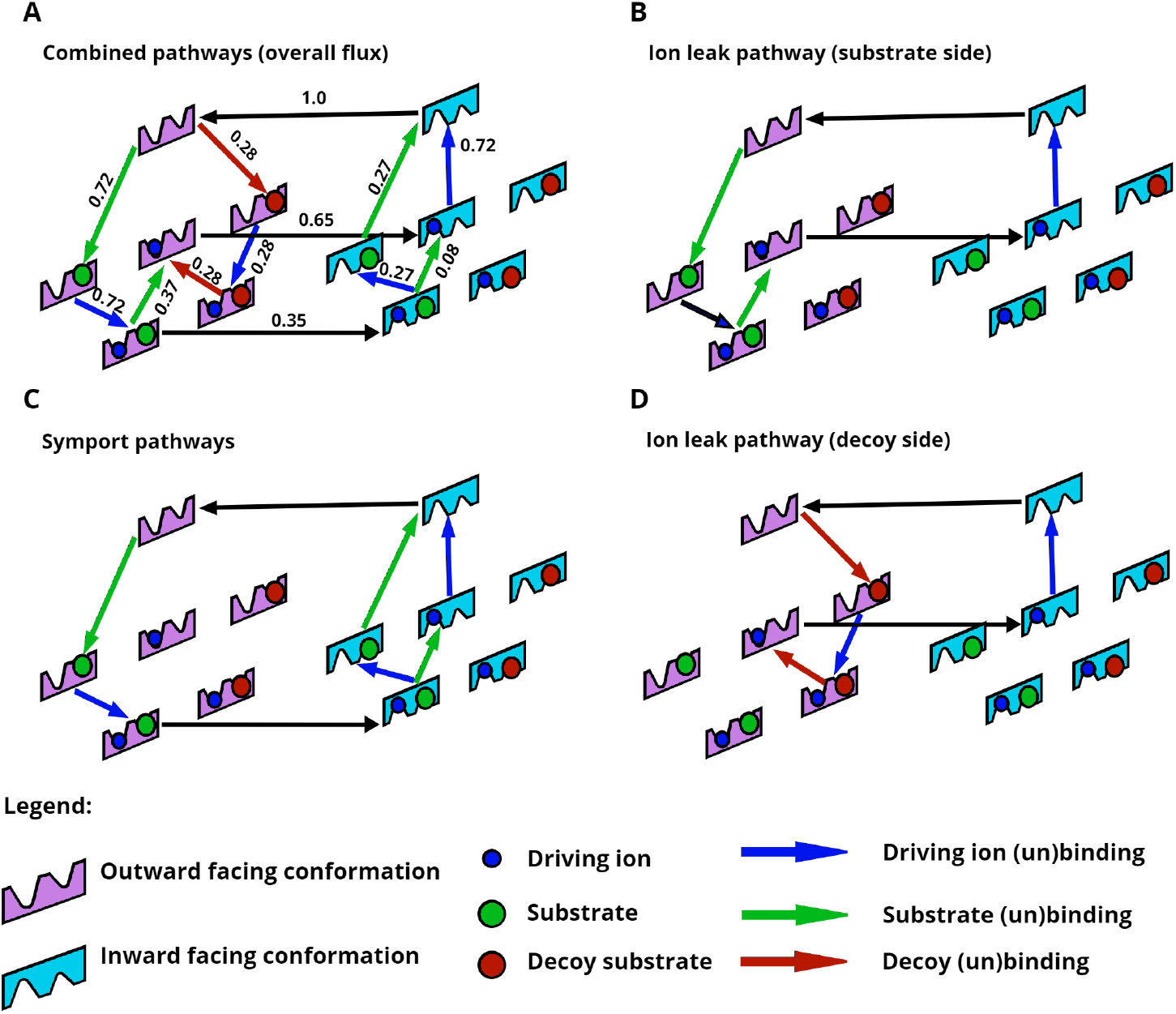
Dissection of a single model exhibiting enhanced selectivity into component pathways. The full model is shown in (**A**) with the net probability flows scaled by the largest edge flow, while panels (B) – (D) are various continuous cycles abstracted from the full model. (**B**) An ion leak pathway in which the substrate and ion bind extracellularly, but only the ion is transported into the cell because the substrate unbinds on the extracellular side. (**C**) A (split) cotransport pathway in which the substrate and ion both are transported into the cell. (**D**) A second ion leak pathway, mirroring (B), in which the decoy and ion bind extracellularly, but only the ion is transported into the cell. Overall, the substrate and decoy are both driven to unbind in the outward-facing conformation, shown on the left, due to ion leak pathways. However, due to the difference in binding affinities between decoy and substrate, the substrate is more likely to rebind and be transported into the intracellular region. The full process employs the ion leaks to enhance selectivity

Selectivity can be examined over a range of driving ion chemical-potential differences, Δ*μ*_ion_ by examining the substrate and decoy substrate fluxes (Fig. 5). The discrimination ratio of substrate to decoy, *J*_substrate_/*J*_decoy_, is essentially perfect at the value used to perform the MC sampling, namely Δ*μ*_ion_ = −4*k_B_T*, where the decoy flux vanishes while the substrate flux remains significant. Interestingly, in the range −4*k_B_T* < Δ*μ*_ion_ ≲ −3*k_B_T*, substrate is pumped into the cell, while the decoy flows out, down its gradient.

**Fig 5.**
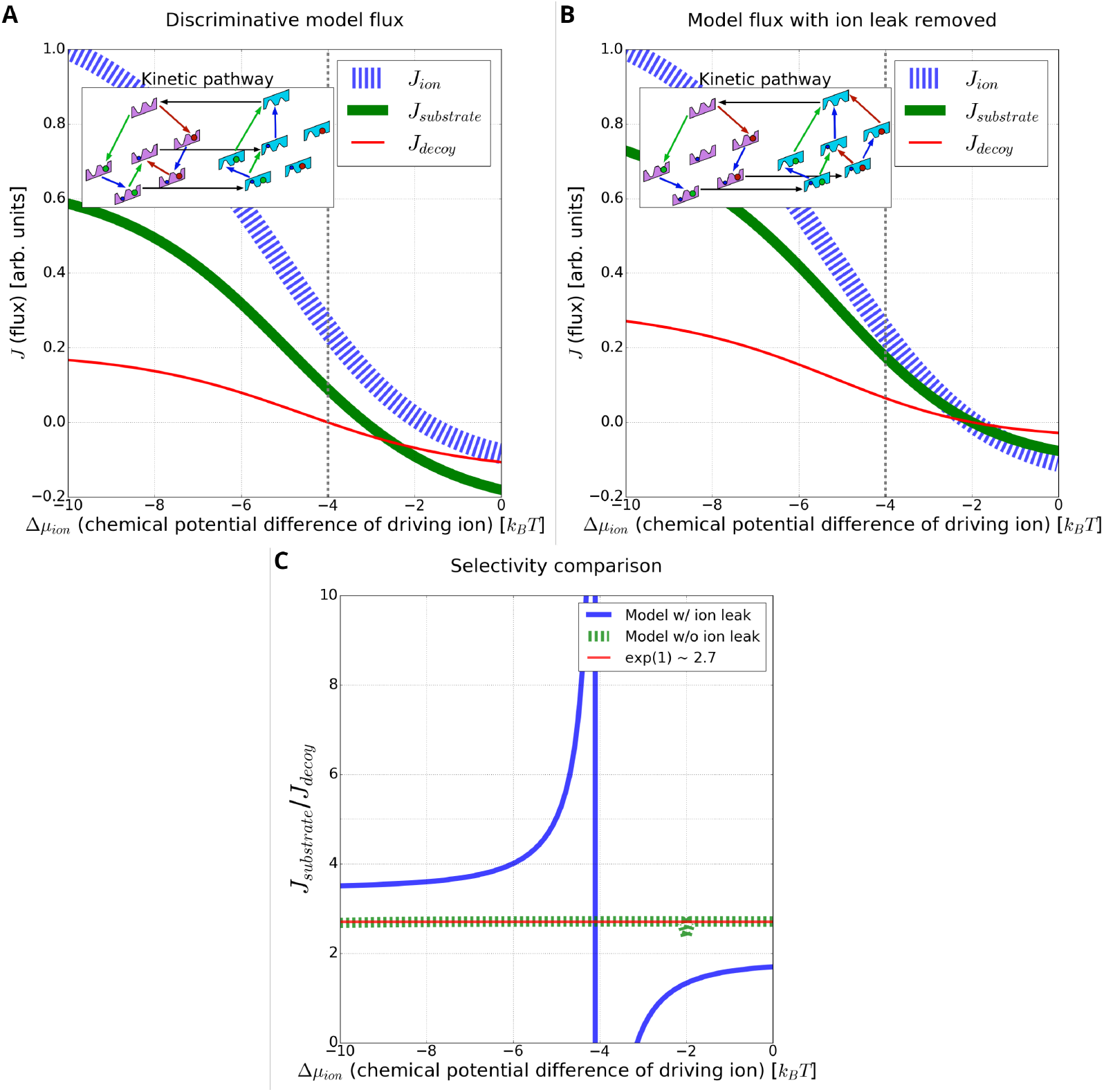
Enhanced discrimination driven by an ion leak. (**A**) For the model of Fig. 4, the substrate, ion, and decoy flux are shown for a varying chemical potential difference of the driving ion. Note the negligible decoy flux relative to the substrate flux near −4*k_B_T*. **Inset** is the kinetic pathway diagram of the model at a specific chemical potential difference (−4*k_B_T*, vertical line) of the ion. (**B**) The same discriminative model as in (A), but with the energy barrier between the ion-only bound states in the inward and outward conformations raised by 100*k_B_T*, effectively shutting off the ion leak. Both the substrate and decoy fluxes increase. **Inset** is the kinetic pathway diagram of the model with the leak removed, resulting in two symmetrical pathways for substrate and decoy transport. (**C**) Comparing the ratio of substrate to decoy flux (selectivity) for the same model with and without an ion leak. With the ion leak, the selectivity approaches infinity due to the negligible decoy flux. In contrast, removing the ion limits the selectivity to the expected equilibrium-like value of *e*^ΔΔ*G*=1^. Note that the sign change of the selectivity is due to the change in substrate and decoy flux direction.

To further investigate the mechanism of enhanced selectivity, we compared the original model B (**Fig. 5A**) to the same model with the ion leak removed (Fig. 5B, note absence of central horizontal flow). The leak-free model – generated by increasing the ion-only-bound conformational transition barrier energy by 100*k_B_T* – has symmetric pathways for both substrate and toxin transport (Fig. 5B inset). Removing the leak dramatically decreases discrimination, as shown in Figure 5C, especially near the ion chemical potential difference Δ*μ*_ion_ = −4*k_B_T* at which MC sampling was performed. In particular, the leak-free model exhibits a decrease in selectivity, approaching the expected equilibrium-like value of *e*^ΔΔ*G*^ ≈ 2.7 with ΔΔ*G* = 1 (see Fig. 5B and **SI-3**) [24]. This suggests that the ion leak is the prime mechanism driving enhanced selectivity for this model.

### Meta-analysis of discriminative models

Our model-sampling approach yields numerous models. We examined them on a “systems basis” using a filtering and clustering procedure.

To start, we filtered for the highest-performing models found during four separate runs of 1e6 MC steps (**see SI-4**). Specifically, models were filtered based on the ion-to-substrate flux ratio being greater than 0.1, and the substrate-to-decoy ratio being greater than 10*e*^ΔΔ*G*^ (ten times the expected discrimination ratio at equilibrium) [24], where ΔΔ*G* = 1 is the difference in binding affinities between the substrate and decoy substrate. These filtering constraints ensured that the analysis only contained models with a sufficient stoichiometry of sodium ions to the substrate, as well as a minimum baseline for enhanced selectivity.

Filtering for performance resulted in 1783 of the aggregate 8000 models exhibiting enhanced selectivity, and we probed these via clustering based on a similarity metric. Clustering revealed four different mechanistic model classes (clusters) with differing kinetic pathways leading to enhanced selectivity (**fig. 6**). Procedurally, for each model in the filtered set, the flows on all edges were scaled by the maximum edge flow in that model. The normalized edge flows define a flow vector used to calculate Euclidean distances and perform complete-linkage clustering [42] based on a distance threshold of 0.65, which we found empirically to reveal qualitatively different pathways.

**Fig 6.**
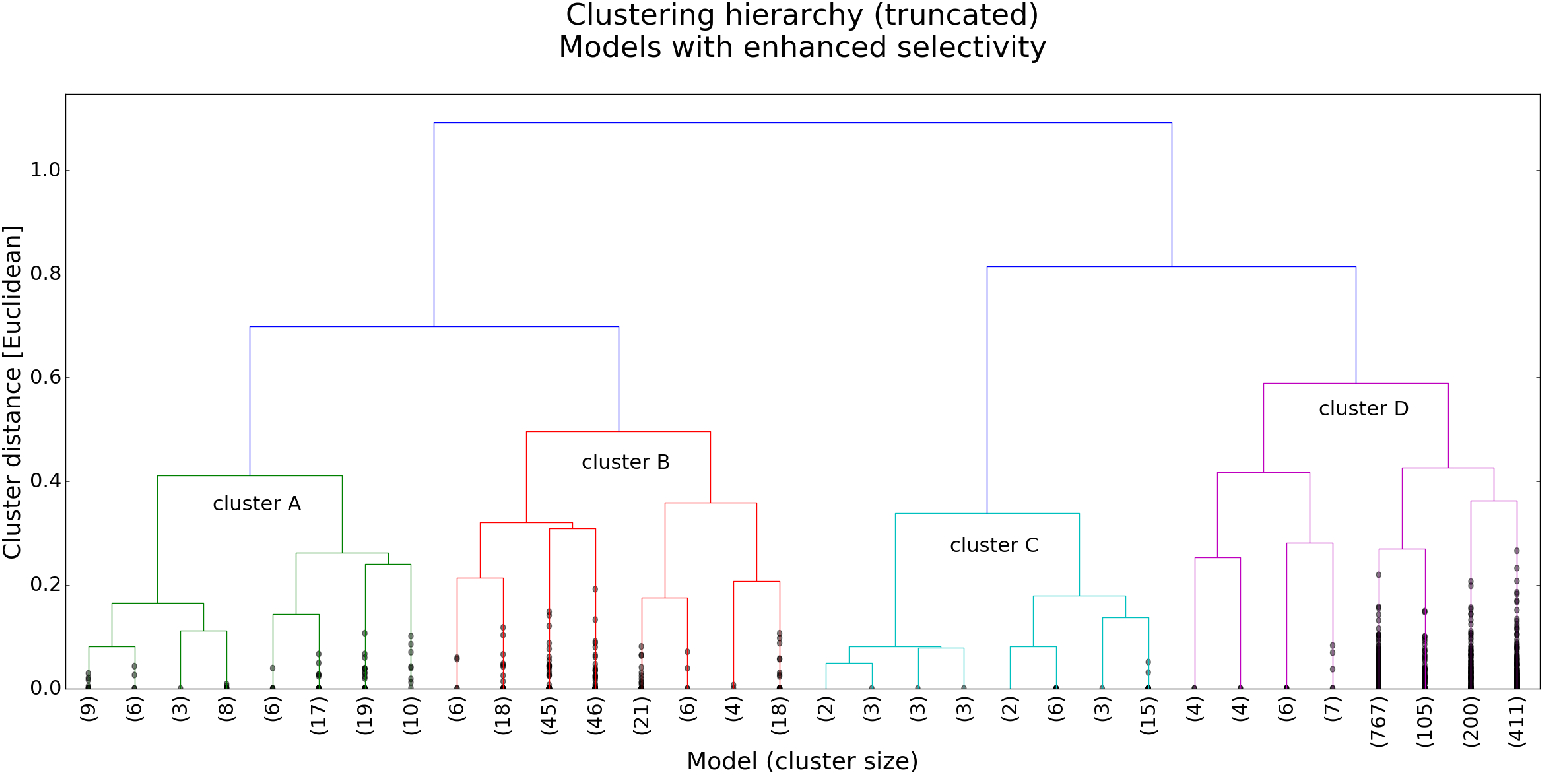
Pathway meta-analysis: Clustering analysis of best-performing models based on model similarity. The dendrogram was truncated to four levels for visual clarity, with the number of models below the truncation shown in parenthesis. Dendrogram created using Python/SciPy and Matplotlib.

All model clusters, which importantly were filtered for enhanced selectivity of substrate over decoy, contain an ion leak. As discussed previously (see Fig. 4), the models shown in **Figure 7** contain “futile” ion-leak cycles, which consist of all the paths that include the central arrow connecting the ion-only-bound outward-facing to inward-facing conformations. In many, but not all models (see models in clusters A, B, D), this ion leak is coupled to substrate and decoy unbinding in the outward-facing state. In other words, the molecules bound to the OF state of the transporter are effectively ejected back to the extracellular space in a process that expends free energy from the ion gradient.

**Fig 7.**
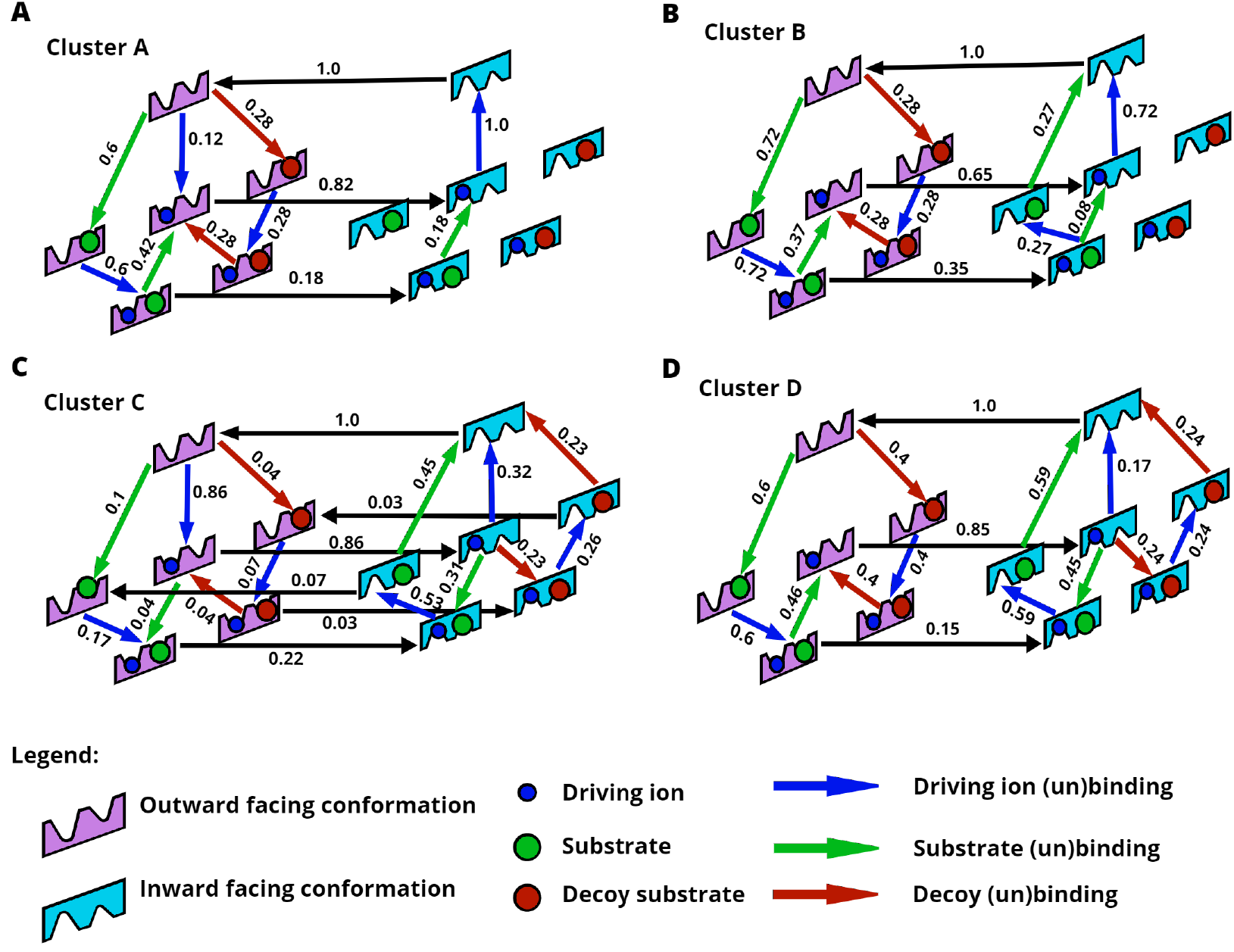
Kinetic pathways of the four model classes (**A-D**) found from clustering analysis (Fig. 6). Each of these models exhibits a high level of selectivity due to an ion leak, as discussed in the text. Note that the net flow values shown along edges are scaled by the maximum flow edge of the individual model.

To further confirm the role of the futile ion cycle in promoting selectivity, the ion leak was removed for each of the representative cluster models (Fig. 7A-D). This was done by raising the corresponding energy barriers as described previously for model B. The resulting leak-free models again exhibited a decrease in selectivity to the expected equilibrium value (*e*^ΔΔ*G*^ with ΔΔ*G* = 1), indicating that the ion leak is indeed the driving mechanism for enhanced selectivity in our models as in Hopfield-like kinetic proofreading [24].

### Model class analysis

It is instructive to examine all the model classes further to understand their differences. Models corresponding to clusters A and B (Fig. 7A,B) share a similar network structure where the substrate (or decoy) tends to bind before the ion, followed by the unbinding of both substrate and decoy, which favors the stronger-binding substrate for transport. A and B differ slightly in that models in cluster A contain an ion-only binding transition in the outward conformation and have a single unbinding pathway for the substrate and ion in the inward-facing conformation (i.e. one symporting pathway). The models in cluster B have two unbinding pathways for the substrate and ion in the inward-facing conformation (i.e. two symporting pathways, see Fig. 4C).

The model class of cluster C (Fig. 7) embodies a much less intuitive mechanism. First, the model is fully connected: each allowed state transition has a non-zero flow. Unlike the other three model classes, class C does not exhibit a net flow of substrate unbinding in the outward-facing conformation. Model C also contains parallel futile cycles with no net transport for either the substrate and decoy in which the ion dissipates its gradient. In the inward-facing conformation, both the substrate and decoy are driven by the ion leak to bind and then unbind again, resulting in more substrate than decoy flux due to their different binding affinities.

Models in cluster D (Fig. 4) contain unbinding steps for both the substrate and decoy in both OF and IF conformations, driven by an ion leak. On the OF side, ion driving appears to force all the decoy to unbind, since there are no horizontal transitions from outward-facing on the left to the corresponding inward-facing states on the left, whereas a fraction of the native substrate remains bound (stronger affinity by ΔΔ*G* = 1*k_B_T*) during the OF-to-IF conformational transition; more precisely, there is an equal and opposite flow of conformational transitions for the decoy-bound states, while the substrate exhibits a net productive flow into the cell. The ion leak also drives the substrate and decoy to bind and unbind in the inward facing conformation, but these processes ‘cancel out’ and do not lead to net flux of either substrate.

For all four models, at low ion chemical potential differences (similar to the magnitude of the substrate/decoy chemical potential difference), there is a regime where the ion and substrate are transported into the cell, and the decoy is transported out of the cell (see Fig. 5A and **SI-3**). This suggests an alternative mode of transport in which the cell ‘detoxifies’ by exporting the decoy substrate.

We can also probe the costs and restrictions on enhanced selectivity. Although our Monte Carlo sampling optimized substrate-to-decoy selectivity at a particular set of chemical-potential differences, the enhanced selectivity generally occurs over a range of thermodynamic driving forces (**see SI-5**). However, some models exhibit high selectivity at a low cost (i.e., low stoichiometric ratio of ion to substrate flux), without any added constraints on the flux ratios. Analysis of the representative models suggests that uniformly high discrimination over a range of thermodynamic conditions occurs with a correspondingly uniformly high cost (see SI), although this is not necessarily a representative sample. This issue will be of interest in future work.

## Discussion

Overall we have seen that Monte Carlo sampling of model space can find diverse, testable models that provide insight into complicated cotransport systems. Unlike prior related work for ion channels [32–34], we have not sought to fit or optimize a single model to a set of experimental data. Rather, we have taken a *systems* approach in asking for *sets of models capable of performing a given function* and applied this to discover multiple transporter models with a discriminatory capability not previously envisioned in the literature, to our knowledge.

First, the model-sampling approach systematically identified ideal and mixture pathways for both simple symporters and antiporters, validating the approach. Subsequent study of a more complex state space including the very realistic possible binding of a “decoy” substrate indicates the possibility of cotransporters that exhibit enhanced selectivity similar to Hopfield’s and Ninio’s kinetic proofreading models [24, 25]. The enhnanced-selectivity models all exhibit an ion leak which, when removed, prevents the enhanced discrimination. This apparently unreported mechanism for secondary active transporters can enhance selectivity to a remarkable degree in a limited range of conditions. Clustering analysis of all models sampled shows a fairly diverse group of model classes that exhibit enhanced selectivity using different kinetic pathways.

Every kinetic model that is generated is also thermodynamically consistent with both equilibrium and non-equlibrium constraints. Since all biochemical reactions are governed by the laws of thermodynamics, this consistency is an important part of accurately modeling the mechanisms of molecular machines. The energy-based formulation of reaction rates *α_ij_*, including non-equilibrium effects where appropriate, combined with an automated state-space construction distinguishes our method from similar approaches.

Although our approach was not developed for the study of evolutionary molecular biology, the method “discovers” working models starting from unproductive models akin to earlier work [30–34]. The changes to rate constants observed in some trajectories may be related to structurally feasible changes. In the future, a more sophisticated approach could attempt to build in additional structural constraints to develop candidate feasible pathways for transporter evolution. We note that the present study already included constraints on the similarity of substrate and decoy, as well as their mutually exclusive binding.

All the enhanced-selectivity models sampled in this study, by construction, were of the ‘Hopfield type’ [24] where substrate discrimination results from the interplay of an external driving force (due to the ion gradient) and a difference in binding affinity between the two substrates. No additional differences between the substrate and decoy were permitted: specifically, all barrier energies were constrained to be identical for both species. This type of proofreading differs from what might be called “internal proofreading” where additional discrimination results from differing barrier heights [26]. Biological proofreading can be expected to mix both mechanisms to some degree [27], but we chose to focus on Hopfield proofreading because an arbitrary degree of discrimination can result from differing barriers – for example, if the decoy species has a negligible on-rate for the transporter. Our approach differs somewhat from Hopfield’s and Nino’s [24, 25] in that no irreversible steps enter our models. The full reversibility appears to be a necessary ingredient in the ideal discrimination (unbounded ratio of substrate to decoy flux) that occurs in some models for specific concentrations.

As emphasized decades ago by Hill [44], in complex networks such as those explored here, it is the network as a whole rather than key steps which define the mechanism. The current networks are simple enough that we can point to intuitive processes coupling ion leaks to unbinding, but the mechanism is defined by the overall process. Undoubtedly, in more realistic networks including a fuller set of conformational states, it will become more difficult to describe an intuitive mechanism. Nevertheless, the principles uncovered in the simple systems can provide useful guidance for conceptualizing complex systems.

## Conclusion

Motivated by evidence for the alternative behavior of molecular machines, we have developed a thermodynamically consistent approach to systematically explore a range of mechanisms, generating multiple experimentally testable kinetic models based on a predetermined fitness function. Prior work has focused on developing individual sets of optimal parameters [32–34] and not on generating model sets, which we believe is essential for developing precise mechanistic hypotheses. The approach was designed for complex state spaces which can be automatically generated, and which would be difficult to analyze by intuition alone. To our knowledge, the platform is the first to enable model sampling building in both equilibrium and non-equilibrium thermodynamic rules. This ‘systems biology’ approach to analyzing mechanisms of molecular machines was applied to a cotransporter state-space with and without a decoy substrate. After validating the method against ‘textbook’ symporter/antiporter models, we generated a variety of mechanisms that enhance selectivity – including Hopfield-like proofreading networks, which could have important biological implications and biotechnology applications. The mechanisms discovered using an automated approach would be difficult to design on an *ad hoc* basis starting from a limited set of experimental structures.

## Supporting information

**S1 Text Detailed methods.**

**S2 Table Simulation parameters.**

**S3.1 Figure Symporter model pathway (without decoy substrate).**

**S3.2 Figure Antiporter model pathway (without decoy substrate).**

**S3.3 Figure Antiporter flux diagram (without decoy substrate.**

**S3.4 Figure Symporter model pathway with ion leak removed (and decoy substrate present).**

**S3.5 Figure Flux diagrams of the representative models for each cluster.**

**S4 Text Model clustering and sampling.**

**S4.1 Figure Trajectory in model space.**

**S4.2 Figure Trajectory in cluster space.**

**S5.1 Figure Cost of the representative models.**

**S5.2 Figure Selectivity of the representative models.**

## Acknowledgments

We thank Barmak Mostofian and Jeremy Copperman for helpful discussions.

## References

1. Alberts B. Molecular biology of the cell. Sixth edition ed. New York, NY: Garland Science, Taylor and Francis Group; 2015.

2. Poulsen SB, Fenton RA, Rieg T. Sodium-glucose cotransport:. Current Opinion in Nephrology and Hypertension. 2015;24(5):463–469. doi:10.1097/MNH.0000000000000152.

3. Demin Jr O, Yakovleva T, Kolobkov D, Demin O. Analysis of the efficacy of SGLT2 inhibitors using semi-mechanistic model. Frontiers in Pharmacology. 2014;5. doi:10.3389/fphar.2014.00218.

4. Deng D, Yan N. GLUT, SGLT, and SWEET: Structural and mechanistic investigations of the glucose transporters: Structural Investigations of the Glucose Transporters. Protein Science. 2016;25(3):546–558. doi:10.1002/pro.2858.

5. Bisignano P, Kalyanaraman C, Ghezzi C, Wright EM, Abramson J, Paz A, et al. Structural Insights into Sodium-Dependent Sugar Transporters and their Inhibition Mechanism. Biophysical Journal. 2017;112(3):128a. doi:10.1016/j.bpj.2016.11.714.

6. Wright EM, Ghezzi C, Loo DDF. Novel and Unexpected Functions of SGLTs. Physiology. 2017;32(6):435–443. doi:10.1152/physiol.00021.2017.

7. Bisignano P, Ghezzi C, Jo H, Polizzi NF, Althoff T, Kalyanaraman C, et al. Inhibitor binding mode and allosteric regulation of Na+-glucose symporters. Nature Communications. 2018;9(1):5245. doi:10.1038/s41467-018-07700-1.

8. Mangold KE, Brumback BD, Angsutararux P, Voelker TL, Zhu W, Kang PW, et al. Mechanisms and models of cardiac sodium channel inactivation. Channels. 2017;11(6):517–533. doi:10.1080/19336950.2017.1369637.

9. Bazzone A, Zabadne AJ, Salisowski A, Madej MG, Fendler K. A Loose Relationship: Incomplete H + /Sugar Coupling in the MFS Sugar Transporter GlcP. Biophysical Journal. 2017;113(12):2736–2749. doi:10.1016/j.bpj.2017.09.038.

10. Henderson RK, Fendler K, Poolman B. Coupling efficiency of secondary active transporters. Current Opinion in Biotechnology. 2019;58:62–71. doi:10.1016/j.copbio.2018.11.005.

11. Kettner C, Bertl A, Obermeyer G, Slayman C, Bihler H. Electrophysiological Analysis of the Yeast V-Type Proton Pump: Variable Coupling Ratio and Proton Shunt. Biophysical Journal. 2003;85(6):3730–3738. doi:10.1016/S0006-3495(03)74789-4.

12. Robinson AE, Thomas NE, Morrison EA, Balthazor BM, Henzler-Wildman KA. New free-exchange model of EmrE transport. Proceedings of the National Academy of Sciences. 2017;114(47):E10083–E10091. doi:10.1073/pnas.1708671114.

13. Shi L, Quick M, Zhao Y, Weinstein H, Javitch JA. The Mechanism of a Neurotransmitter:Sodium Symporter—Inward Release of Na+ and Substrate Is Triggered by Substrate in a Second Binding Site. Molecular Cell. 2008;30(6):667–677. doi:10.1016/j.molcel.2008.05.008.

14. Hill TL. Free Energy Transduction and Biochemical Cycle Kinetics. New York, NY: Springer New York; 1989. Available from: http://link.springer.com/10.1007/978-1-4612-3558-3.

15. Schnoerr D, Sanguinetti G, Grima R. Approximation and inference methods for stochastic biochemical kinetics—a tutorial review. Journal of Physics A: Mathematical and Theoretical. 2017;50(9):093001. doi:10.1088/1751-8121/aa54d9.

16. Goutsias J, Jenkinson G. Markovian dynamics on complex reaction networks. Physics Reports. 2013;529(2):199–264. doi:10.1016/j.physrep.2013.03.004.

17. Galstyan V, Funk L, Einav T, Phillips R. Combinatorial Control through Allostery. The Journal of Physical Chemistry B. 2019;123(13):2792–2800. doi:10.1021/acs.jpcb.8b12517.

18. Lim W, Lee C, Tang C. Design Principles of Regulatory Networks: Searching for the Molecular Algorithms of the Cell. Molecular Cell. 2013;49(2):202–212. doi:10.1016/j.molcel.2012.12.020.

19. Sobczak I, Lolkema JS. The 2-Hydroxycarboxylate Transporter Family: Physiology, Structure, and Mechanism. Microbiology and Molecular Biology Reviews. 2005;69(4):665–695. doi:10.1128/MMBR.69.4.665-695.2005.

20. Berg JM, Tymoczko JL, Stryer L, Stryer L. Biochemistry. 5th ed. New York: W.H. Freeman; 2002.

21. Mitchell P. A General Theory of Membrane Transport From Studies of Bacteria. Nature. 1957;180(4577):134–136. doi:10.1038/180134a0.

22. Jardetzky O. Simple Allosteric Model for Membrane Pumps. Nature. 1966;211(5052):969–970. doi:10.1038/211969a0.

23. Zuckerman, Daniel M. Physical Lens on the Cell: Advanced Cycle Logic; 2014. Available from: https://www.physicallensonthecell.org/chemical-physics/advanced-cycle-logic.

24. Hopfield JJ. Kinetic Proofreading: A New Mechanism for Reducing Errors in Biosynthetic Processes Requiring High Specificity. Proceedings of the National Academy of Sciences. 1974;71(10):4135–4139. doi:10.1073/pnas.71.10.4135.

25. Ninio J. Kinetic amplification of enzyme discrimination. Biochimie. 1975;57(5):587–595. doi:10.1016/S0300-9084(75)80139-8.

26. Fersht AR. Editing mechanisms in protein synthesis. Rejection of valine by the isoleucyl-tRNA synthetase. Biochemistry. 1977;16(5):1025–1030. doi:10.1021/bi00624a034.

27. Banerjee K, Kolomeisky AB, Igoshin OA. Elucidating interplay of speed and accuracy in biological error correction. Proceedings of the National Academy of Sciences. 2017;114(20):5183–5188. doi:10.1073/pnas.1614838114.

28. Mallory JD, Kolomeisky AB, Igoshin OA. Trade-Offs between Error, Speed, Noise, and Energy Dissipation in Biological Processes with Proofreading. The Journal of Physical Chemistry B. 2019;123(22):4718–4725. doi:10.1021/acs.jpcb.9b03757.

29. Murugan A, Huse DA, Leibler S. Discriminatory Proofreading Regimes in Nonequilibrium Systems. Physical Review X. 2014;4(2):021016. doi:10.1103/PhysRevX.4.021016.

30. Deckard A, Sauro HM. Preliminary Studies on the In Silico Evolution of Biochemical Networks. ChemBioChem. 2004;5(10):1423–1431. doi:10.1002/cbic.200400178.

31. Gottstein W, Müller S, Herzel H, Steuer R. Elucidating the adaptation and temporal coordination of metabolic pathways using in-silico evolution. Biosystems. 2014;117:68–76. doi:10.1016/j.biosystems.2013.12.006.

32. Gurkiewicz M, Korngreen A. A numerical approach to ion channel modelling using whole-cell voltage-clamp recordings and a genetic algorithm. PLoS computational biology. 2007;3(8):e169.

33. Menon V, Spruston N, Kath WL. A state-mutating genetic algorithm to design ion-channel models. Proceedings of the National Academy of Sciences. 2009;106(39):16829–16834. doi:10.1073/pnas.0903766106.

34. Teed ZR, Silva JR. A computationally efficient algorithm for fitting ion channel parameters. MethodsX. 2016;3:577–588. doi:10.1016/j.mex.2016.11.001.

35. Ollivier JF, Shahrezaei V, Swain PS. Scalable Rule-Based Modelling of Allosteric Proteins and Biochemical Networks. PLoS Computational Biology. 2010;6(11):e1000975. doi:10.1371/journal.pcbi.1000975.

36. C Mason J, W Covert M. An energetic reformulation of kinetic rate laws enables scalable parameter estimation for biochemical networks. Journal of Theoretical Biology. 2019;461:145–156. doi:10.1016/j.jtbi.2018.10.041.

37. Sekar JAP, Hogg JS, Faeder JR. Energy-based modeling in BioNetGen. In: 2016 IEEE International Conference on Bioinformatics and Biomedicine (BIBM). Shenzhen, China: IEEE; 2016. p. 1460–1467. Available from: http://ieeexplore.ieee.org/document/7822739/.

38. Metropolis N, Rosenbluth AW, Rosenbluth MN, Teller AH, Teller E. Equation of State Calculations by Fast Computing Machines. The Journal of Chemical Physics. 1953;21(6):1087–1092. doi:10.1063/1.1699114.

39. Hastings WK. Monte Carlo sampling methods using Markov chains and their applications. Biometrika. 1970;57(1):97–109. doi:10.1093/biomet/57.1.97.

40. Phillips R, Kondev J, Theriot J. Physical biology of the cell. New York: Garland Science; 2009.

41. Whitelam S, Geissler PL. Avoiding unphysical kinetic traps in Monte Carlo simulations of strongly attractive particles. The Journal of Chemical Physics. 2007;127(15):154101. doi:10.1063/1.2790421.

42. Rokach L, Maimon O. Clustering Methods. In: Maimon O, Rokach L, editors. Data Mining and Knowledge Discovery Handbook. New York: Springer-Verlag; 2005. p. 321–352. Available from: http://link.springer.com/10.1007/0-387-25465-X_15.

43. Adelman JL, Ghezzi C, Bisignano P, Loo DDF, Choe S, Abramson J, et al. Stochastic steps in secondary active sugar transport. Proceedings of the National Academy of Sciences. 2016;113(27):E3960–E3966. doi:10.1073/pnas.1525378113.

44. Hill TL. Free energy transduction in biology: the steady-state kinetic and thermodynamic formalism. New York: Academic Press; 1977.

