## Supplementary material for "A systems-biology approach to molecular machines: Exploration of alternative transporter mechanisms": SI

### 1 Detailed methods

#### 1.1 State name definitions

| | |
|---|---|
| OF | Outward-facing conformation |
| IF | Inward-facing conformation |
| No | Extracellular sodium ion |
| Ni | Intracellular sodium ion |
| Nb | Bound sodium ion |
| So | Extracellular substrate |
| Si | Intracellular substrate |
| Sb | Bound substrate |
| Wo | Extracellular decoy substrate |
| Wi | Intracellular decoy substrate |
| Wb | Bound decoy substrate |

#### 1.2 Equivalent states and transitions

We have defined groups of states that are physically equivalent at steady state with fixed extracellular and intracellular concentrations. After a species (e.g. ion or substrate) is transported, the physical state will remain the same because of the steady-state assumption. As an example, consider a hypothetical transporter of substrate (S) driven by a sodium ion (N). The state 'OF-Nb-So' describes the extracellular (outward) facing conformation (OF) with sodium-bound (Nb) and substrate unbound and in the extracellular region (So). This is physically equivalent to the state 'OF-Nb-Si', which only differs by the "location" of the unbound substrate (Si, substrate inside the cell). The location of a substrate or ion is needed in order to fully identify transitions – i.e., the origin (inside or outside) of substrate or ion in a binding process.

Transitions are similarly grouped based on states that are equivalent under the conditions stated above. Considering the same hypothetical transporter: an extracellular-to-bound sodium transition (No $\rightarrow$ Nb) would be physically equivalent for extracellular (So) and intracellular (Si) substrate in the outward-facing conformation (OF). The equivalent state and transition groups are constrained to share the same state or transition energy during the Monte Carlo (MC) energy perturbations for self-consistency.

To investigate the Hopfield kinetic proofreading model of transport, we have an additional constraint that groups 'equivalent' transitions for the substrate and decoy substrate. As an example, an inward-to-outward facing conformational transition with only the decoy bound (e.g. OF-No-So-Wb to IF-No-So-Wb) would be equivalent to an inward-to-outward facing conformational transition with only the substrate bound (e.g. OF-No-Sb-Wo to IF-No-Sb-Wo). This extra constraint prohibits a difference in transition energies between equivalent substrate and decoy transitions; effectively removing "internal proofreading" models from our search. See

##### 1.2.1 List of equivalent states/transitions for a cotransporter without decoy substrate

Equivalent states:

- IF-Ni-So, IF-No-Si, IF-No-So, IF-Ni-Si
- OF-No-So, OF-Ni-Si, OF-No-Si, OF-Ni-So
- IF-Ni-Sb, IF-No-Sb
- IF-Nb-So, IF-Nb-Si
- OF-No-Sb, OF-Ni-Sb
- OF-Nb-So, OF-Nb-Si

Equivalent transitions:

- OF-Ni-Sb⟷OF-Ni-Si, OF-No-Sb⟷OF-No-Si
- OF-Ni-So⟷OF-Ni-Sb, OF-No-So⟷OF-No-Sb
- IF-Nb-Si⟷IF-Ni-Si, IF-Nb-So⟷IF-Ni-So
- OF-No-So⟷OF-Nb-So, OF-No-Si⟷OF-Nb-Si
- OF-No-So⟷IF-No-So, OF-Ni-Si⟷IF-Ni-Si, OF-Ni-So⟷IF-Ni-So, OF-No-Si⟷IF-No-Si
- OF-Ni-Sb⟷IF-Ni-Sb, OF-No-Sb⟷IF-No-Sb
- IF-No-So⟷IF-Nb-So, IF-No-Si⟷IF-Nb-Si,
- OF-Nb-Si⟷OF-Ni-Si, OF-Nb-So⟷OF-Ni-So,
- OF-Nb-So⟷IF-Nb-So, OF-Nb-Si⟷IF-Nb-Si
- IF-No-So⟷IF-No-Sb, IF-Ni-So⟷IF-Ni-Sb,
- IF-No-Sb⟷IF-No-Si, IF-Ni-Sb⟷IF-Ni-Si

##### 1.2.2 List of equivalent states/transitions for a cotransporter with decoy substrate

Equivalent states:

- OF-Nb-So-Wb, OF-Nb-Sb-Wo, OF-Nb-Sb-Wi, OF-Nb-Si-Wb
- IF-Ni-So-Wb, IF-No-Sb-Wo, IF-Ni-Sb-Wo, IF-No-Sb-Wi, IF-Ni-Si-Wb, IF-No-Si-Wb, IF-Ni-Sb-Wi, IF-No-So-Wb
- OF-No-So-Wb, OF-No-Si-Wb, OF-Ni-So-Wb, OF-Ni-Sb-Wo, OF-No-Sb-Wo, OF-Ni-Si-Wb, OF-Ni-Sb-Wi, OF-No-Sb-Wi
- IF-Nb-So-Wo, IF-Nb-Si-Wo, IF-Nb-Si-Wi, IF-Nb-So-Wi
- IF-Ni-So-Wo, IF-Ni-Si-Wo, IF-No-Si-Wi, IF-No-So-Wo, IF-No-Si-Wo, IF-Ni-So-Wo, IF-Ni-So-Wi, IF-No-So-Wi, IF-Ni-Si-Wi
- IF-Nb-So-Wb, IF-Nb-Sb-Wi, IF-Nb-Si-Wb, IF-Nb-Sb-Wo
- OF-Nb-So-Wo, OF-Nb-Si-Wo, OF-Nb-Si-Wi, OF-Nb-So-Wi
- OF-No-So-Wo, OF-Ni-So-Wi, OF-No-Si-Wi, OF-Ni-Si-Wo, OF-Ni-Si-Wi, OF-No-Si-Wo, OF-No-So-Wi, OF-Ni-So-Wo

Equivalent transitions:

- IF-No-So-Wb$\longleftrightarrow$IF-Nb-So-Wb, IF-No-Sb-Wi$\longleftrightarrow$IF-Nb-Sb-Wi, IF-No-Sb-Wo$\longleftrightarrow$IF-Nb-Sb-Wo, IF-No-Si-Wb$\longleftrightarrow$IF-Nb-Si-Wb
- OF-Nb-So-Wi$\longleftrightarrow$IF-Nb-So-Wi, OF-Nb-Si-Wo$\longleftrightarrow$IF-Nb-Si-Wo, OF-Nb-Si-Wi$\longleftrightarrow$IF-Nb-Si-Wi, OF-Nb-So-Wo$\longleftrightarrow$IF-Nb-So-Wo
- IF-Nb-So-Wb$\longleftrightarrow$IF-Nb-So-Wi, IF-Nb-Si-Wb$\longleftrightarrow$IF-Nb-Si-Wi, IF-Nb-Sb-Wo$\longleftrightarrow$IF-Nb-Si-Wo, IF-Nb-Sb-Wi$\longleftrightarrow$IF-Nb-Si-Wi
- OF-Nb-Si-Wb$\longleftrightarrow$IF-Nb-Si-Wb, OF-Nb-So-Wb$\longleftrightarrow$IF-Nb-So-Wb, OF-Nb-Sb-Wo$\longleftrightarrow$IF-Nb-Sb-Wo, OF-Nb-Sb-Wi$\longleftrightarrow$IF-Nb-Sb-Wi
- IF-Nb-Sb-Wi$\longleftrightarrow$IF-Ni-Sb-Wi, IF-Nb-So-Wb$\longleftrightarrow$IF-Ni-So-Wb, IF-Nb-Si-Wb$\longleftrightarrow$IF-Ni-Si-Wb, IF-Nb-Sb-Wo$\longleftrightarrow$IF-Ni-Sb-Wo
- OF-No-Sb-Wo$\longleftrightarrow$OF-No-Si-Wo, OF-No-Sb-Wi$\longleftrightarrow$OF-No-Si-Wi, OF-Ni-Si-Wb$\longleftrightarrow$OF-Ni-Si-Wi, OF-No-Si-Wb$\longleftrightarrow$OF-No-Si-Wi, OF-Ni-Sb-Wo$\longleftrightarrow$OF-Ni-Si-Wo, OF-Ni-So-Wb$\longleftrightarrow$OF-Ni-So-Wi, OF-No-So-Wb$\longleftrightarrow$OF-No-So-Wi, OF-Ni-Sb-Wi$\longleftrightarrow$OF-Ni-Si-Wi
- OF-Ni-Sb-Wo$\longleftrightarrow$IF-Ni-Sb-Wo, OF-Ni-So-Wb$\longleftrightarrow$IF-Ni-So-Wb, OF-No-Sb-Wi$\longleftrightarrow$IF-No-Sb-Wi, OF-Ni-Sb-Wi$\longleftrightarrow$IF-Ni-Sb-Wi, OF-No-Sb-Wo$\longleftrightarrow$IF-No-Sb-Wo, OF-Ni-Si-Wb$\longleftrightarrow$IF-Ni-Si-Wb, OF-No-Si-Wb$\longleftrightarrow$IF-No-Si-Wb, OF-No-So-Wb$\longleftrightarrow$IF-No-So-Wb
- IF-No-Si-Wb$\longleftrightarrow$IF-No-Si-Wi, IF-Ni-Sb-Wo$\longleftrightarrow$IF-Ni-Si-Wo, IF-No-So-Wb$\longleftrightarrow$IF-No-So-Wi, IF-No-Sb-Wi$\longleftrightarrow$IF-No-Si-Wi, IF-Ni-So-Wb$\longleftrightarrow$IF-Ni-So-Wi, IF-Ni-Sb-Wi$\longleftrightarrow$IF-Ni-Si-Wi, IF-No-Sb-Wo$\longleftrightarrow$IF-No-Si-Wo, IF-Ni-Si-Wb$\longleftrightarrow$IF-Ni-Si-Wi
- OF-Ni-Si-Wi$\longleftrightarrow$IF-Ni-Si-Wi, OF-No-So-Wi$\longleftrightarrow$IF-No-So-Wi, OF-Ni-Si-Wo$\longleftrightarrow$IF-Ni-Si-Wo, OF-No-So-Wo$\longleftrightarrow$IF-No-So-Wo, OF-Ni-So-Wo$\longleftrightarrow$IF-Ni-So-Wo, OF-No-Si-Wo$\longleftrightarrow$IF-No-Si-Wo, OF-No-Si-Wi$\longleftrightarrow$IF-No-Si-Wi, OF-Ni-So-Wi$\longleftrightarrow$IF-Ni-So-Wi

- OF-Nb-Sb-Wo⟷OF-Nb-Si-Wo, OF-Nb-So-Wb⟷OF-Nb-So-Wi, OF-Nb-Sb-Wi⟷OF-Nb-Si-Wi, OF-Nb-Si-Wb⟷OF-Nb-Si-Wi
- IF-No-Si-Wi⟷IF-Nb-Si-Wi, IF-No-Si-Wo⟷IF-Nb-Si-Wo, IF-No-So-Wi⟷IF-Nb-So-Wi, IF-No-So-Wo⟷IF-Nb-So-Wo
- OF-Nb-So-Wi⟷OF-Nb-Sb-Wi, OF-Nb-Si-Wo⟷OF-Nb-Si-Wb, OF-Nb-So-Wo⟷OF-Nb-So-Wb, OF-Nb-So-Wo⟷OF-Nb-Sb-Wo
- IF-Nb-So-Wo⟷IF-Ni-So-Wo, IF-Nb-Si-Wi⟷IF-Ni-Si-Wi, IF-Nb-Si-Wo⟷IF-Ni-Si-Wo, IF-Nb-So-Wi⟷IF-Ni-So-Wi
- OF-No-Si-Wi⟷OF-Nb-Si-Wi, OF-No-Si-Wo⟷OF-Nb-Si-Wo, OF-No-So-Wi⟷OF-Nb-So-Wi, OF-No-So-Wo⟷OF-Nb-So-Wo
- OF-Nb-Si-Wb⟷OF-Ni-Si-Wb, OF-Nb-So-Wb⟷OF-Ni-So-Wb, OF-Nb-Sb-Wo⟷OF-Ni-Sb-Wo, OF-Nb-Sb-Wi⟷OF-Ni-Sb-Wi
- OF-No-Sb-Wo⟷OF-Nb-Sb-Wo, OF-No-So-Wb⟷OF-Nb-So-Wb, OF-No-Si-Wb⟷OF-Nb-Si-Wb, OF-No-Sb-Wi⟷OF-Nb-Sb-Wi
- IF-No-Si-Wo⟷IF-No-Si-Wb, IF-Ni-Si-Wo⟷IF-Ni-Si-Wb, IF-Ni-So-Wi⟷IF-Ni-Sb-Wi, IF-Ni-So-Wo⟷IF-Ni-Sb-Wo, IF-No-So-Wo⟷IF-No-So-Wb, IF-No-So-Wo⟷IF-No-Sb-Wo, IF-Ni-So-Wo⟷IF-Ni-So-Wb, IF-No-So-Wi⟷IF-No-Sb-Wi
- OF-Nb-So-Wi⟷OF-Ni-So-Wi, OF-Nb-Si-Wo⟷OF-Ni-Si-Wo, OF-Nb-Si-Wi⟷OF-Ni-Si-Wi, OF-Nb-So-Wo⟷OF-Ni-So-Wo
- OF-No-So-Wi⟷OF-No-Sb-Wi, OF-No-So-Wo⟷OF-No-So-Wb, OF-Ni-So-Wo⟷OF-Ni-Sb-Wo, OF-No-Si-Wo⟷OF-No-Si-Wb, OF-No-So-Wo⟷OF-No-Sb-Wo, OF-Ni-Si-Wo⟷OF-Ni-Si-Wb, OF-Ni-So-Wo⟷OF-Ni-So-Wb, OF-Ni-So-Wi⟷OF-Ni-Sb-Wi
- IF-Nb-Si-Wo⟷IF-Nb-Si-Wb, IF-Nb-So-Wo⟷IF-Nb-Sb-Wo, IF-Nb-So-Wo⟷IF-Nb-So-Wb, IF-Nb-So-Wi⟷IF-Nb-Sb-Wi

#### 1.3 Tempering

##### 1.3.1 Automated tempering

Automated tempering tracks the change in fitness of the previous models, decreasing $\beta$ (heating) if the fitness has not changed over several models, and then increasing $\beta$ (cooling) once a user-defined threshold is met. An initial $\beta$ is set and then checked at fixed Monte Carlo step intervals. At these intervals, the fractional Monte Carlo energy difference from that previous checkpoint is calculated: $E_{MC}^{\text{frac}} = 2\frac{E_{MC}^{\text{new}} - E_{MC}^{\text{old}}}{|E_{MC}^{\text{new}}| - |E_{MC}^{\text{old}}|}$ where $E_{MC}^{\text{new}}$ is the current Monte Carlo energy and $E_{MC}^{\text{old}}$ is the Monte Carlo energy at the previous checkpoint. If the fractional energy is approximately constant, $|E_{MC}^{frac}| <$ tolerance, $\beta$ decreases (heats) by a user constrained scale factor. If the fractional energy difference has decreased, $\beta$ increases (cools) by a user constrained scale factor, subject to a user-defined probability to stay at the current beta, $P_{\beta}^{\text{stay}}$. If the fractional energy has increased, $\beta$ decreases (heats) by a user constrained scale factor, subject to a user-defined probability to stay at the current beta, $P_{\beta}^{\text{stay}}$.

##### 1.3.2 Manual tempering

Manual tempering uses a fixed schedule that adjusts $\beta$ by a set amount at each Monte Carlo step interval. The user defines the minimum and maximum $\beta$ (inverse temperature), and also defines how many Monte Carlo steps to remain at the minimum and maximum $\beta$, decrease $\beta$ from maximum to minimum, and to increase $\beta$ from minimum to maximum. This schedule can then be scaled by a user-defined factor for further sampling optimization. For enhanced selectivity simulations, the default tempering schedule (found empirically) is: 125 MC steps at the minimum $\beta$, 100 MC steps increasing $\beta$, 1450 MC steps at the maximum $\beta$, and 325 MC steps decreasing $\beta$.

#### 1.4 Flux calculation

The flux of a given species x, $J_x$, is calculated by summing the net flows along a user-defined set of transitions for that species. For the ion, this is the set of transitions from an ion-bound state to an ion-inside state (Nb$\rightarrow$Ni). The substrate flux is calculated from the set of transitions from a substrate-bound state to a substrate-inside state (Sb$\rightarrow$Si). Similarly, decoy flux is calculated from the set of transitions from a decoy-bound state to a decoy-inside state (Wb$\rightarrow$Wi). For simplicity, we have removed 'backdoor' transport which would allow a species to be transported in the opposing direction of the conformational state (e.g. OF-Nb$\rightarrow$OF-Ni). Due these added constraints, the flux for a given species is calculated using only the net flows of the (un)binding transitions for that species in the *inward-facing conformation.*

### 2 Simulation parameters

#### 2.1 Cotransporter without decoy substrate

| | **Symporter** | **Antiporter** |
|---|---|---|
| MC steps | 1e6 | 1e6 |
| Random seed | 123456 | 123456 |
| Maximum $\Delta E$ for state/transition $[k_B T]$ | 1.0 | 1.0 |
| Tempering schedule | Automatic | Automatic |
| Tempering tolerance | 0.3 | 0.3 |
| $\beta$ initial $[k_B T]^{-1}$ | 1e1 | 1e1 |
| $\beta$ scale factor | 1e3 | 1e3 |
| $P_\beta^{\text{stay}}$ | 0.2 | 0.2 |
| MC steps to change $\beta$ | 2e2 | 2e2 |
| $\Delta\mu_{\text{ion}}$ $[k_B T]$ | -4 | -4 |
| $\Delta\mu_{\text{substrate}}$ $[k_B T]$ | +2 | -2 |
| Rate prefactor, $k_0$ [s$^{-1}$] | 1e-3 | 1e-3 |
| Energy function, $E_{MC}$ | $-J_{\text{substrate}}$ | $+J_{\text{substrate}}$ |

#### 2.2 Cotransporter with decoy substrate

| | **Run 1** | **Run 2** | **Run 3** | **Run 4** |
|---|---|---|---|---|
| MC steps | 1e6 | 1e6 | 1e6 | 1e6 |
| Random seed | 456789 | 456789 | 456789 | 456789 |
| Maximum $\Delta E$ for state/transition $[k_B T]$ | 1.0 | 0.5 | 0.2 | 1.0 |
| Tempering schedule | Manual | Manual | Manual | Manual |
| Tempering scale factor | 1.0 | 0.5 | 1.0 | 2.0 |
| $\beta_{\min}$ $[k_B T]^{-1}$ | 1e-100 | 1e-100 | 1e-100 | 1e-100 |
| $\beta_{\max}$ $[k_B T]^{-1}$ | 1e30 | 1e30 | 1e30 | 1e30 |
| MC steps to change $\beta$ | 1 | 1 | 1 | 1 |
| $\Delta\mu_{\text{ion}}$ $[k_B T]$ | -4 | -4 | -4 | -4 |
| $\Delta\mu_{\text{substrate}}$ $[k_B T]$ | 2 | 2 | 2 | 2 |
| $\Delta\mu_{\text{decoy}}$ $[k_B T]$ | 2 | 2 | 2 | 2 |
| $\Delta\Delta G$ $[k_B T]$ | 1 | 1 | 1 | 1 |
| Rate prefactor, $k_0$ [s$^{-1}$] | 1e-3 | 1e-3 | 1e-3 | 1e-3 |
| Energy function, $E_{MC}$ | $-J_{\text{substrate}} \frac{\lvert J_{\text{substrate}}\rvert+\epsilon}{\lvert J_{\text{decoy}}\rvert+\epsilon}$ | ... | ... | ... |
| Numerical stability constant, $\epsilon$ | 1e-15 | 1e-15 | 1e-15 | 1e-15 |

### 3 Additional model pathway and flux diagrams

#### 3.1 Symporter model pathway (without decoy substrate).

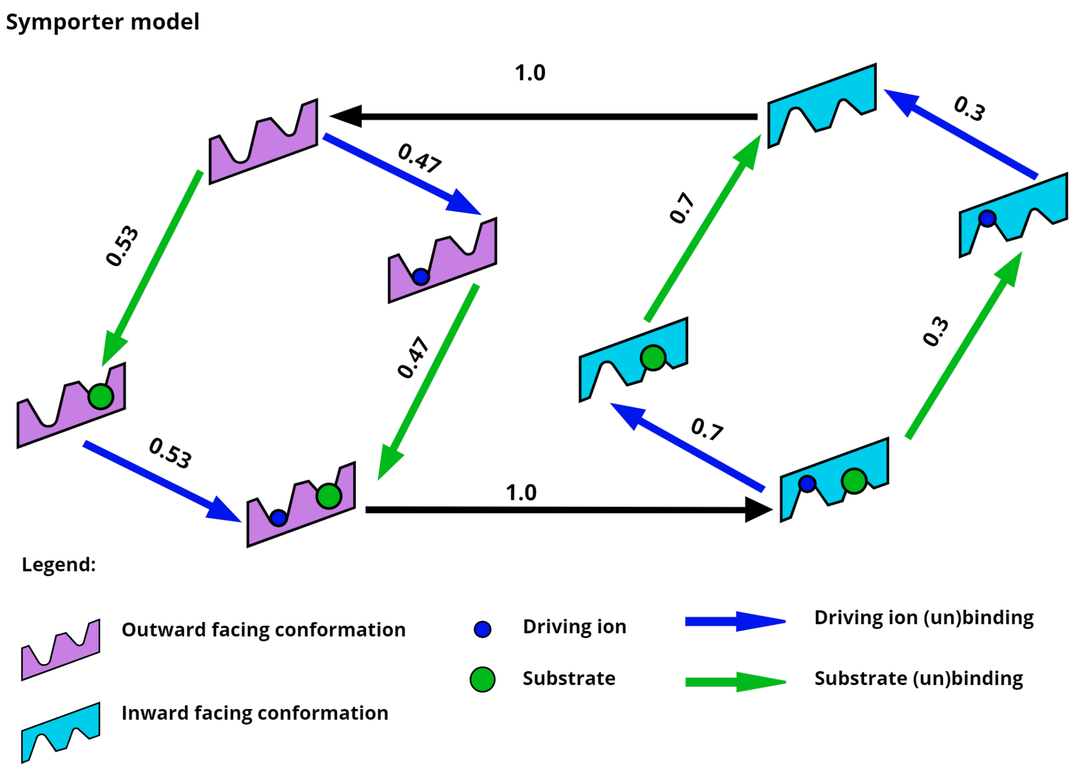


**Figure S3.1:** Pathway of a symporter model found at MC step = 1800. This model exhibits a combination of all four ideal symporter pathways which result in the intracellular transport of one substrate per ion. Note that the flows are scaled by the largest flow edge.

#### 3.2 Antiporter model pathway (without decoy substrate).

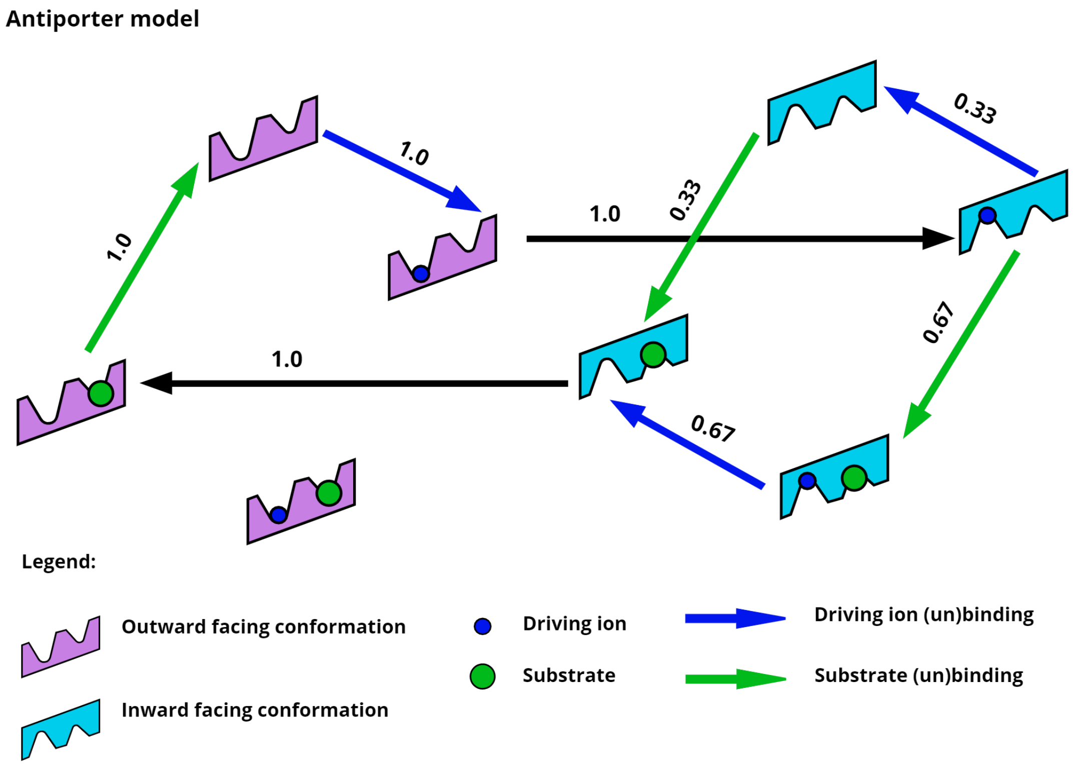


**Figure S3.2:** Pathway of an anitporter model found during the antiporter simulation run at MC step = 845000. This model exhibits a combination of two ideal antiporter pathways which result in the extracellular transport of one substrate per ion. Note that the flows are scaled by the largest flow edge

#### 3.3 Antiporter flux diagram (without decoy substrate).

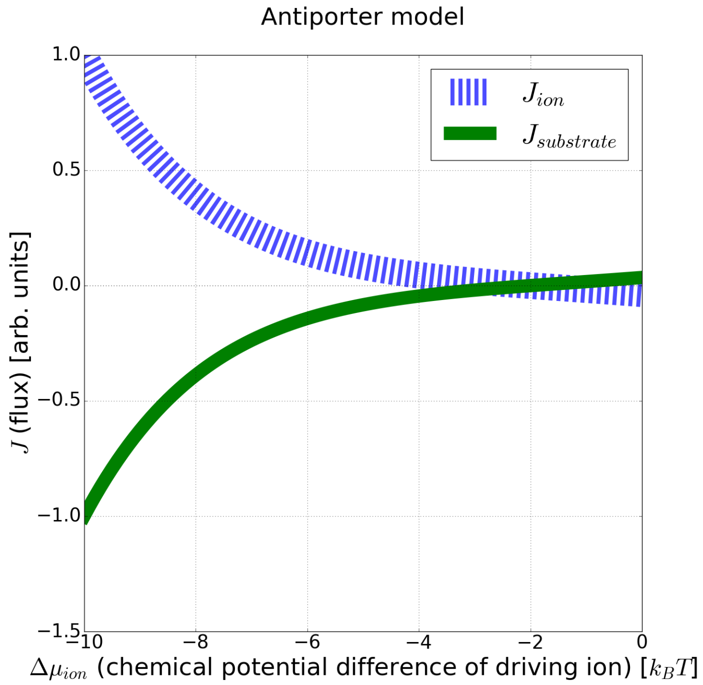


**Figure S3.3:** Flux of an antiporter model found during the antiporter simulation run at MC step = 845000, analyzed over a range of ion chemical potential differences. This model exhibits a 1:1 ratio of ion influx to substrate efflux over a wide range of ion chemical potential differences. Note that the fluxes are scaled by the largest flux value.

#### 3.4 Symporter model with ion leak removed (and with decoy substrate present)

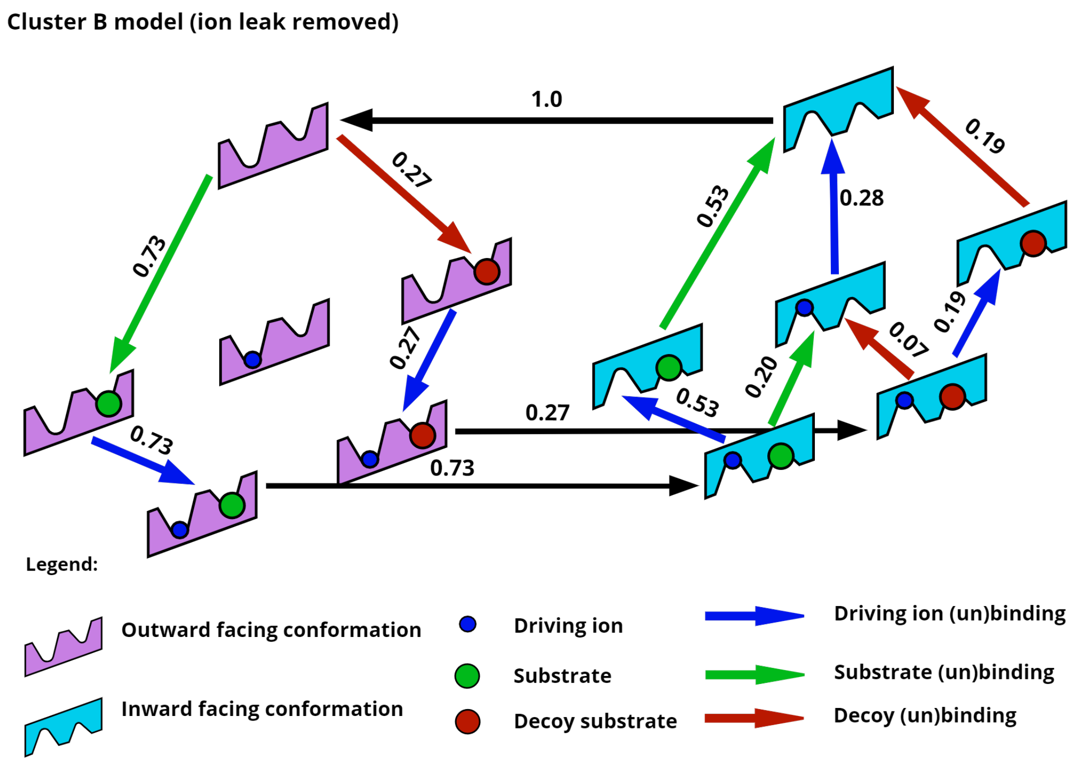


**Figure S3.4:** Pathway of the model with enhanced selectivity representing cluster B, with the futile ion cycle removed. The energy barrier between the ion-only bound states in the inward and outward conformations was raised by 100 $k_BT$, effectively shutting off the ion leak. Note the two symmetrical pathways for substrate and decoy transport. The net flows have been scaled by the maximum flow edge.

#### 3.5 Flux diagrams of the representative models for each cluster

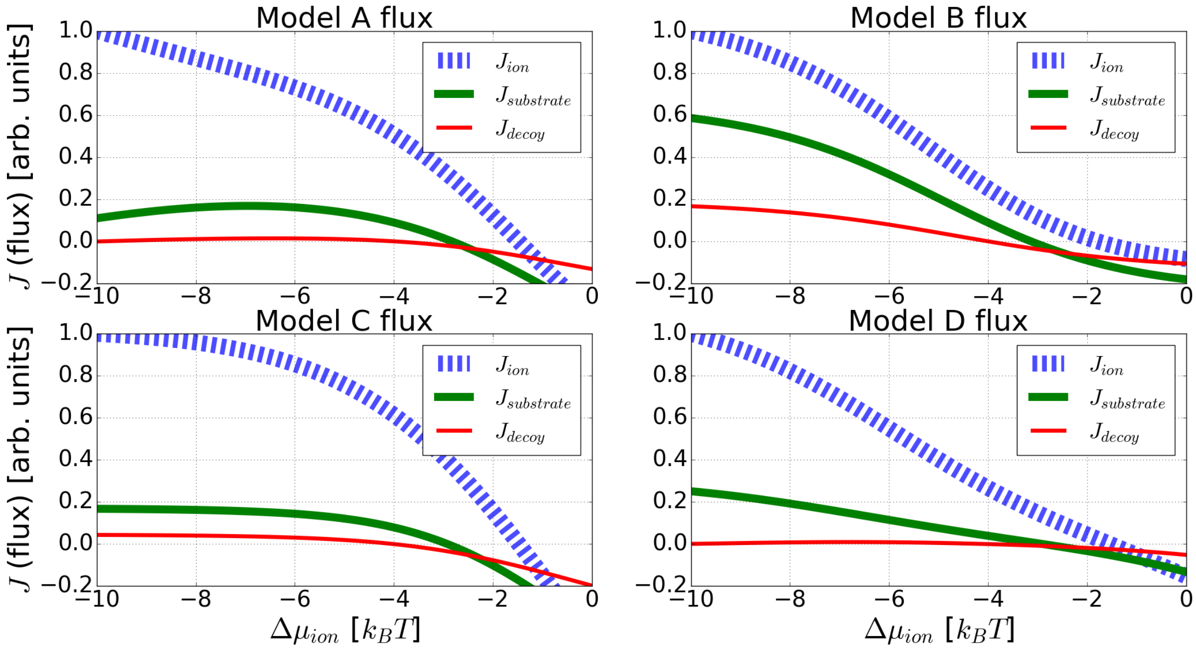


**Figure S3.5:** Flux of the representative model for each cluster, scaled by the maximum flux, over a range of ion chemical potential differences. Each model has a narrow regime where the toxin flows down its gradient out of the cell, while the substrate is driven into the cell by the ion. Near the optimized conditions for the simulation, $\Delta\mu_{ion} = -4k_BT$, these models have negligible decoy flux, resulting in an unbounded substrate to decoy discrimination ratio.

### 4 Model clustering and sampling

Simulations were run for transporters in a 'competitive' environment with a decoy. Clusters were determined using hierarchical clustering with complete-linkage and the Euclidean distance between the scaled flows of each model. The threshold of 0.65 was determined empirically to produce qualitatively different kinetic pathways. This method produced four separate clusters. Representative models corresponding to each cluster were analyzed in the main text and SI.

The representative models used:

| | Run | MC step number |
|---|---|---|
| Cluster A model | 1 | 3500 |
| Cluster B model | 1 | 29000 |
| Cluster C model | 3 | 3000 |
| Cluster D model | 3 | 829000 |

#### 4.1 Trajectory in model space

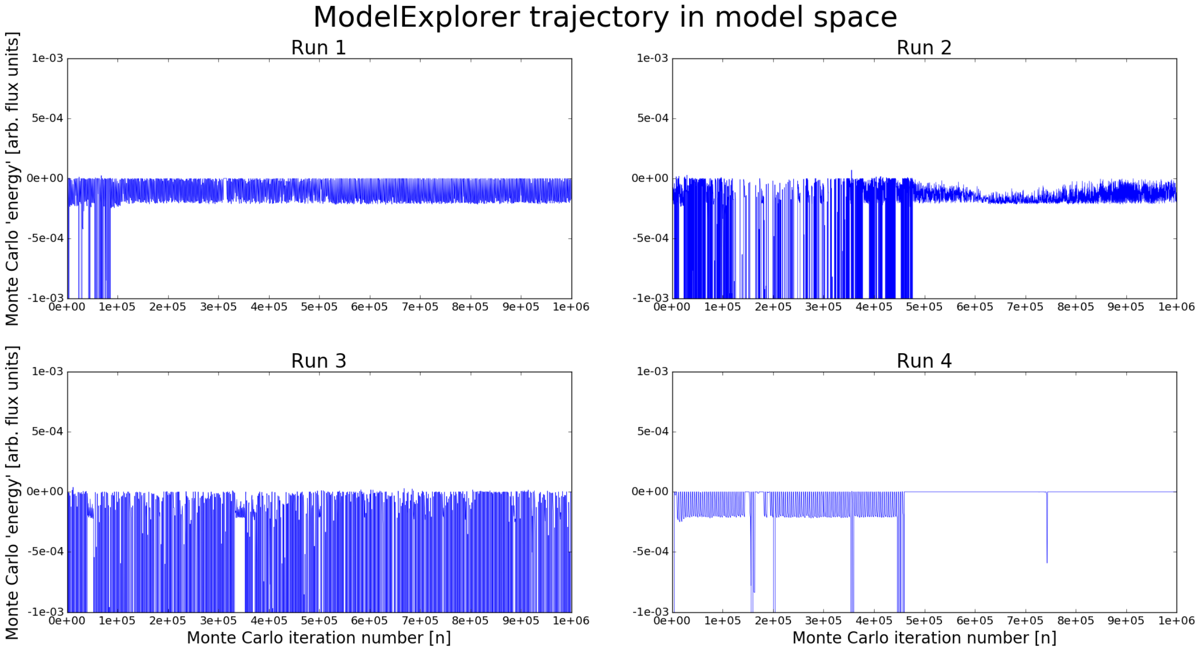


**Figure S4.1:** Trajectory in model space of four different 1e6 Monte Carlo (MC) step simulations at different sampling settings. Simulations were run for transporters in a 'competitive' environment with a decoy using MC energy function: $-J_{\text{substrate}}\frac{|J_{\text{substrate}}|+\epsilon}{|J_{\text{decoy}}|+\epsilon}$ where $J_{\text{substrate}}$ and $J_{\text{decoy}}$ are the fluxes of the substrate and decoy respectively, and $\epsilon$ = 1e-15. Lower MC energy values denote models that are more fit, by convention. As shown in the figures, the tempering schedule aids in avoiding low energy basins. Note that each point on the trajectory is a kinetic model.

#### 4.2 Trajectory in cluster space

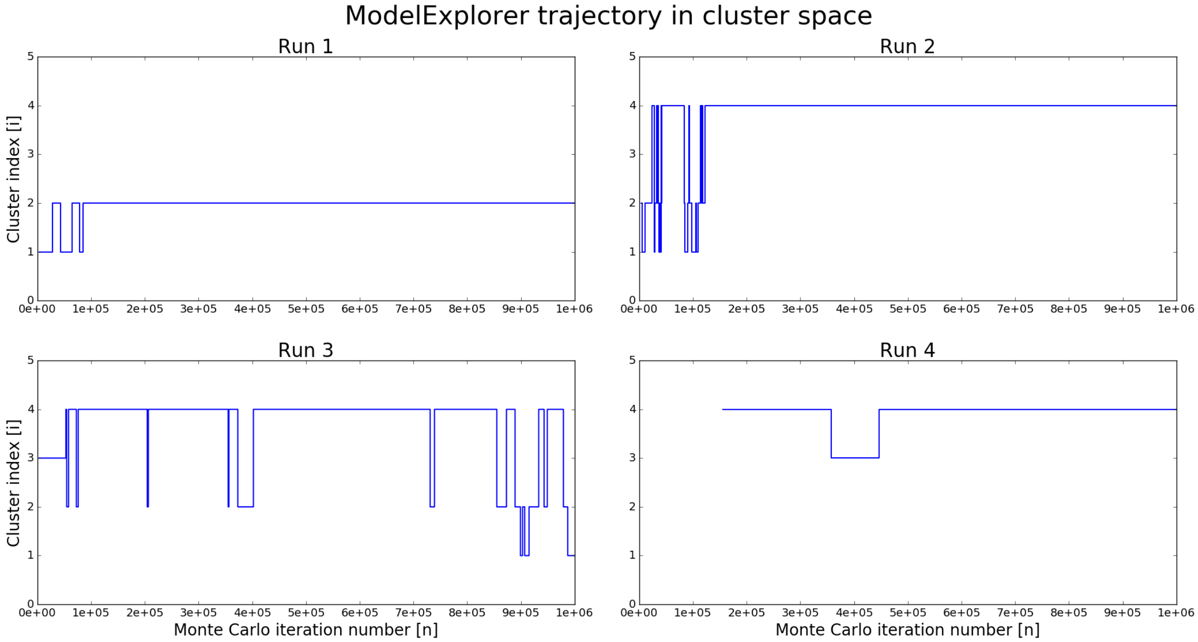


**Figure S4.2:** Trajectory in cluster space of four different 1e6 MC step simulations. Simulations were run for transporters in a 'competitive' environment with a decoy. Models were filtered based on to models with a cost (ion to substrate flux ratio) below 10, and selectivity (substrate to decoy ratio) above $10e^{\Delta\Delta G=1}$. Clusters were determined using hierarchical clustering with complete-linkage and the Euclidean distance between the scaled flows of each model. The threshold of 0.65 was determined empirically to produce qualitatively different kinetic pathways. These graphs indicate that each run only finds a few model classes during the simulation – implying the need for improved sampling methods. Note that in run 4, models meeting the selection criteria (i.e. cost and selectivity) were not found until approximately 1.5e5 MC steps.

### 5 Cost and selectivity analysis

#### 5.1 Cost of representative models

Here we examine the ion to substrate stoichiometry (cost) for each model cluster.

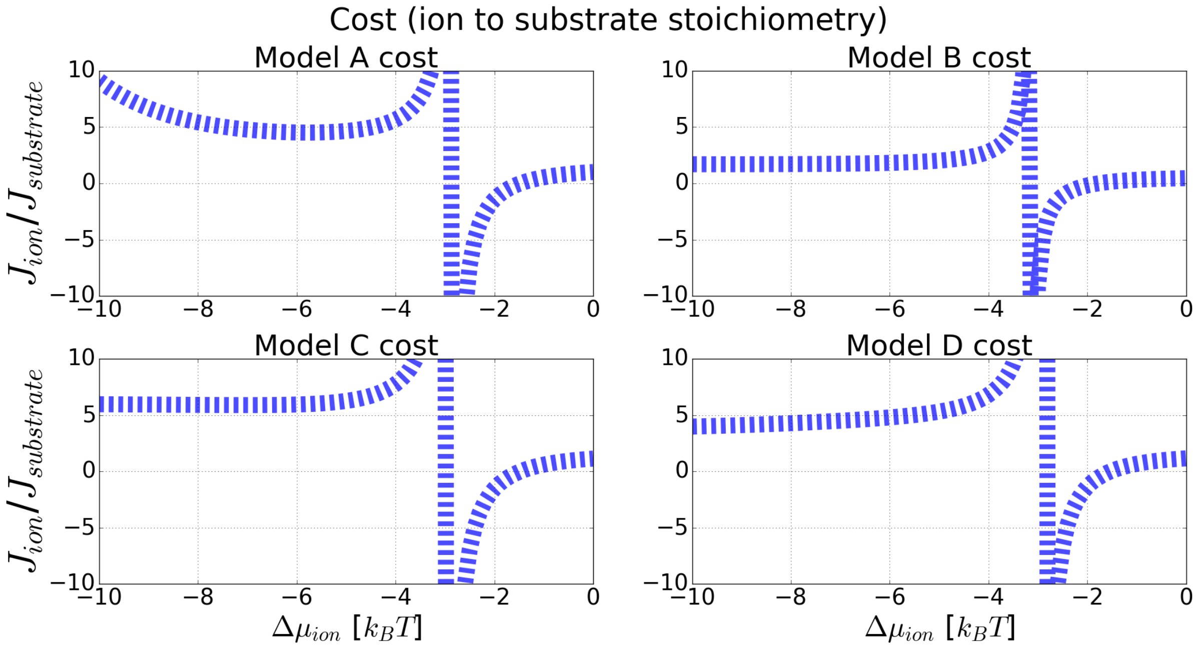


**Figure S5.1:** Cost of the representative model for each cluster, over a range of ion chemical potential differences. All of these models exhibit a cost above the ideal 1:1 stoichiometric ratio for a wide range of chemical potential differences. This extra ions transported relative to the substrate suggest a futile ion transport cycle - i.e. an ion leak. Note that the cost was not included as a constraint in the energy function

#### 5.2 Selectivity of representative models

Here we examine the substrate to decoy stoichiometry (selectivity) for each model cluster.

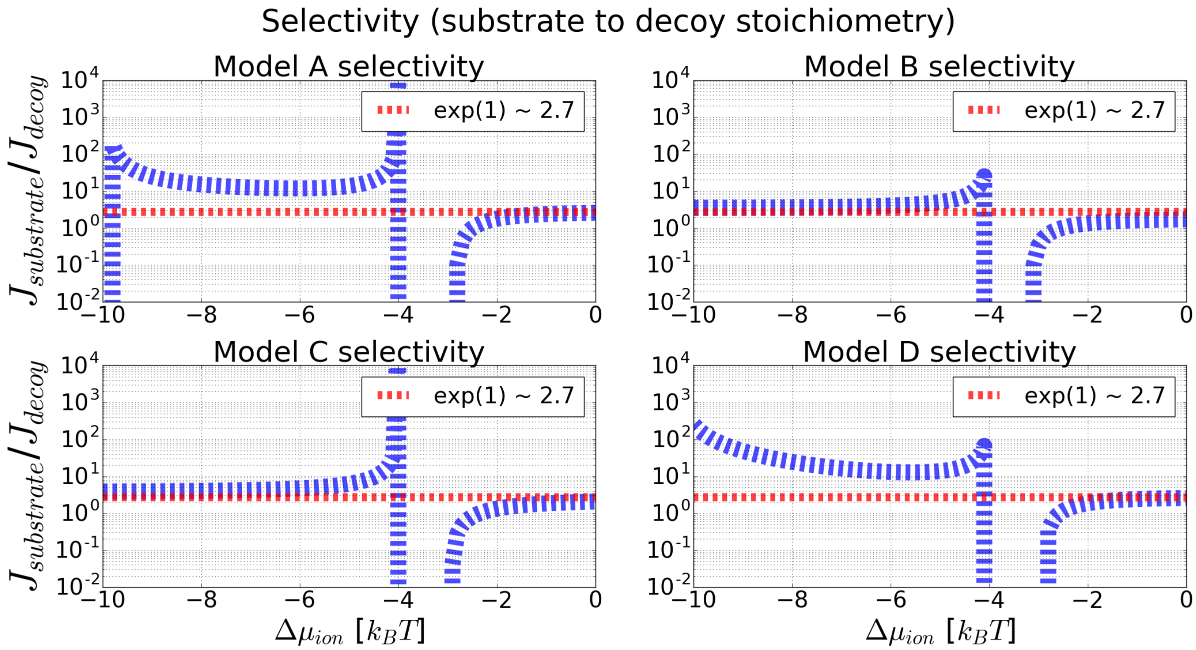


**Figure S5.2:** The stoichiometric ratio of the substrate to ion flux (selectivity), over a range of ion chemical potential differences. All the models demonstrate enhanced selectivity over a range of chemical potential differences, and unbounded selectivity at the optimized condition (at $\Delta\mu_{ion} = 4k_BT$ ). Models A and D exhibit enhanced discrimination over a wide range of conditions. The expected equilibrium value ($e^{\Delta\Delta G=1}$) is shown as a reference. Note that this stoichiometric ratio was used as a primary constraint in our energy function.
